# Characterization of *Plasmodium falciparum* RAP domain proteins-RAP291 and RAP070, and their association with ribosomal RNAs

**DOI:** 10.1101/2022.09.21.508831

**Authors:** Asif Akhtar, Arunaditya Deshmukh, Ashutosh Panda, Ekta Saini, Rajan Pandey, Iqbal Taliy Junaid, Noah Machuki Onchieku, Sadaf Parveen, Suneet Shekhar Singh, Dinesh Gupta, Michael Theisen, Asif Mohmmed, Pawan Malhotra

## Abstract

*Plasmodium* genomes encode multiple RAP (RNA-binding domain abundant in Apicomplexan) domain proteins that contain a conserved module of 56 to 73 amino acids. Here, we characterized two of the *P. falciparum* RAP domain proteins; PfRAP291 & PfRAP070, for their expression and role at asexual blood stages. RNA binding assays and high-throughput CLIP-seq analysis showed that these proteins mainly bind ribosome associated RNAs. Blue-native PAGE and protein-protein interaction studies suggested association of these proteins with MSP-1 complex. Anti-PfRAP291 and anti-PfRAP070 antibodies showed moderate inhibitions in *in-vitro* merozoite invasion assays. Together, these results suggest multiple roles of these proteins; PfRAP291 and PfRAP070, in merozoite invasion and in ribosome regulation during asexual stages of the parasite.

## Introduction

The *Plasmodium* life cycle involves multiple stages with different morphologies in human host as well as in mosquito vector. Each stage requires a well-organized developmental program with specific regulation of gene expression and protein synthesis [1,2]. Although parasite genome encodes low number of transcriptional regulators, however post-transcriptional regulations have been shown to play important role(s) in parasite protein expression regulations [3,4]. Furthermore, comparative studies on parasite’s transcriptomics and proteomics have revealed that translational regulations also play critical role(s) in parasite life cycle [5–7]. For example, during intraerythrocytic development half-life of many mRNAs are extended during the schizont stage, and also many mRNAs are kept in translationally repressed state during gametocyte and schizont stages as they are needed in subsequent stages of development [8]. *Plasmodium* structural RNAs and mRNAs are thus extensively regulated and RNA binding proteins (RBPs) having RNA binding domains (RBDs) are important players in such regulations [9,10]. In human at least 1500 RBPs have been identified so far among which RNA recognition motifs (RRMs) alone are most abundant, constituting more than 200 RBPs [11–13]. RRMs, Zinc-finger domains, K homology domain (KH), Pumilio and Fem3 binding factor (Puf), Acetylation lower binding affinity family (Alba) and RNA helicases are among the best characterized RBPs in human [12–16]. In *Plasmodium*, a recent bioinformatic study has identified 189 RBPs including 72 with RRM, 11 with KLH, 2 with Puf domain, 6 with Alba domain, 31 with Zinc finger domain and 48 having helicase domains [9]. One of the abundant RBD identified in apicomplexans, particularly in *Plasmodium* spp., is RAP domain and its role in apicomplexan has not been yet characterized fully [17–19].

RNA-binding domain abundant in apicomplexans (RAP) is a ~60-residues domain first identified in human MGC5297 protein and is particularly abundant in apicomplexans [17,18]. Although the biological significance of the RAP domain proteins in eukaryotes, in particular among apicomplexans, is yet to be established, nevertheless presence of RAP domain in proteins such as *C. reinhardtii* Raa3 that binds to tscRNA as a part of Ribonucleoprotein complex and Fas-activated serine/threonine kinase (FASTK) that interacts with TIA-1, a downstream effector of eIF2 pathway predict RNA binding function for RAP domains [20,21]. Recently, two *Plasmodium* RAP proteins have been validated as mitochondrial rRNAs binder and have been suggested to play role(s) in mitoribosome regulation [22].

We have previously reported a MSP-1 complex consisting of 11 merozoite surface proteins; MSP-1, MSP-3, MSP-6, MSP-7, MSP-9, RhopH3, RhopH1, RAP-1, RAP-2 and two uncharacterized RAP domain proteins; PfRAP291 (Pf3D7_1029800) & PfRAP070 (Pf3D7_0815100) by immuno-pulldown experiments with merozoite lysate using anti-MSP-1 and anti-PfRhopH3 antibodies [23]. Presence of the two RAP domain proteins; PfRAP291 and PfRAP070, on merozoite surface was intriguing and required further authentication. The present study has been designed to characterize these two *P. falciparum* RAP domain proteins, PF3D7_1029800 (PfRAP291) and PF3D7_0815100 (PfRAP070), for their roles in RNA binding, if any, and also for their expression at asexual blood stages. *In vitro* crosslinking studies followed by immunoprecipitation suggested that these proteins bind parasite ribosomal RNA(s) and are also a part of a MSP-1 complex, suggesting their possible role(s) in invasion as well as in ribosome assembly and/or function.

## Methods

### *In vitro Plasmodium falciparum* culture

*Plasmodium falciparum* strain 3D7 was cultured on human erythrocytes in RPMI-1640 media (Invitrogen) with 4% haematocrit supplemented with 0.5% AlbuMAX® I (gibco). Parasite cultures were maintained using standard protocol described by Trager and Jensen [24]. Synchronization of parasite cultures were carried out using two consecutive sorbitol treatments four hours apart [25].

### Genome screening for RAP domain proteins in *Plasmodium* genomes

The amino acid sequences of RAP proteins were retrieved from PlasmoDB (v49). Its physicochemical properties were identified using ProtParam (Expasy). Conserved domains were identified using Conserved Domain Database (CDD), the Simple Modular Architecture Research Tool (SMART) and Protein Family Database (PFAM). The full deduced amino acid sequence and individual conserved domains were subjected to BLAST (BLASTP) to identify orthologs in PlasmoDB and NCBI protein database. Next, we performed OrthoMCL (version 5) database search to determine the orthologs of *P. falciparum*. A multiple sequence alignment was performed using the retrieved sequences, using T-COFFEE version-11 with default settings. These aligned sequences were further put to analysis by MEME motif search tool for the presence of common residues. The phylogenetic tree was inferred using the Neighbor-Joining method for computing the evolutionary distance with default setting in Molecular Evolutionary Genetics Analysis software (MEGA 7.0). Gaps and missing data were treated using partial deletion method with 95% site-coverage cut-off and 1000 bootstrap replicate to generate phylogenetic tree.

### Cloning of PfRAP291 & PfRAP070 protein fragments and their expression

PfRAP291 & PfRAP070 gene fragments; 465bp for PfRAP291 and 837bp for PfRAP070 were amplified from the genomic DNA of *Plasmodium falciparum 3D7* using the primer pairs; PfRAP291 Forward Primer-5’-CGCCATGGAAATGTTTATTTGTTCAAGACCTCAGCA-3’, PfRAP291 Reverse Primer-5’-CGCCTAGGCTCGAGTACATGAATTTGCATTTGTTGT TTATTATTTTC-3’, PfRAP070 Forward Primer-5’-GCCCATGGAAATGCCACATAA AGATTATTTAGGTGA-3’, PfRAP070 Reverse Primer-5’-GCCCTAGGCTCGAGGGC ATTATGTCCATTTGAGC-3’. The PCR products were cloned into pJET vector (Promega) and sequenced. The fragments were subcloned in the pET28b expression vector between NcoI and XhoI sites. The pET28b constructs of PfRAP291 & PfRAP070 were transformed into Shuffle-30 (NEB) expression host cells. Each culture was induced at 0.5 mM IPTG for 10 h at 37 °C. The cells were disrupted by sonication in lysis buffer (0.05 M Tris, pH 8.0, 0.15 M NaCl, 0.01 M DTT, 1 mM PMSF, 1% Triton X-100) with 9 s pulses at 9 s intervals for 10 times using mini probe. The soluble and insoluble fractions were separated by centrifugation and analysed by SDS-PAGE followed by Western-blot analysis using anti-His antibody. The two recombinant RAP proteins (rPfRAP070 and rPfRAP291) were expressed in soluble form in *E. Coli* shuffle cells. Both the RAP proteins were purified from the supernatant using affinity-based Ni-NTA^+^(nitrilotriacetic acid) chromatography and eluted fractions containing >90% pure proteins were pooled and dialysed against 0.05 M Tris (pH 8), 0.015 M NaCl and 10% glycerol. Finally, the purified recombinant proteins, rPfRAP070 and rRAP291, were stored at −80 °C in aliquots.

### Generation of antibodies against rPfRAP291 & rPfRAP070 proteins

Antibodies against rPfRAP291 & rPfRAP070 were raised in both mice and rabbit. The animals were housed and handled in accordance with the institutional and national guidelines. The institutional animal ethical committee at ICGEB, New Delhi, India, approved the animal use protocol described in the studies. The animals were bred under the guidelines of the authorizing committee. For the antibody generation, five to six weeks old female BALB/c mice were immunized with 25 μg of each protein: rPfRAP291 and rPfRAP070 emulsified in Freund’s complete adjuvant on day 0, followed by three boosters of proteins emulsified with Freund’s incomplete adjuvant on days 14, 28 and 42. The animals were bled for serum collection on day 49. For rabbit immunization, New Zealand white female rabbits were immunized with 200 μg of either of the following recombinant proteins: rPfRAP291 & rPfRAP070 emulsified in Freund’s complete adjuvant on day 0, followed by three boosters emulsified with Freund’s incomplete adjuvant on days 21, 42 and 63. The animals were bled for serum collection on day 70. Antibody titres in serum samples were quantified by enzyme-linked immunosorbent assay (ELISA). Production of antibodies against rPfMSP165, rPfMSP3N and rPfRhopH3b, have been described earlier [26–28].

### Western blot analysis of *Plasmodium falciparum* 3D7 merozoite

Briefly, *P. falciparum* merozoites were harvested as described by Hill et al., 2014 [29] and lysed with equal volumes of RIPA buffer for 1 hour on ice. Then, the merozoite suspensions was triturated several times with 1 ml syringe attached to 26-gauge needle. High speed centrifugation at 15,000 × g for 20 min was carried out to remove insoluble material and the parasite lysate was run on SDS-PAGE and transferred to nitrocellulose membrane. The membranes were probed with rabbit anti-PfRAP291 (1:10,000) or anti-PfRAP070 (1:10,000) antisera followed by goat anti-rabbit HRP conjugated secondary antibody (1:50,000). The membranes were developed with ECL reagent (Bio-Rad) and imaged with Chemi-Doc (Bio-Rad).

### *In vivo* RNA binding analysis by UV-Crosslinking of *P. falciparum* culture and isolation of bound RNA proteins

Briefly, parasite cultures in late schizont stage (44-48 hpi) with 10% parasitemia were harvested, washed with PBS, and then resuspended in cold PBS. For crosslinking with media, the parasites were briefly centrifuged and excess of media were removed leaving enough media for suspension to get monolayer of cells. The resuspended cells were transferred to 10 cm tissue culture dish and placed on ice. The parasites were then irradiated with 254 nm UV light to a total energy of 600 mJ/cm^2^ with intermittent mixing on ice [30–32]. The UV crosslinked cells were harvested by centrifugation.

After cross-linking, parasite infected red blood cells were mixed and homogenised in TRIzol^TM^ (Invitrogen) and the homogenized lysate was incubated at room temperature (RT) for 5 min to dissociate any unstable RNA–protein interactions. The interphase purification was carried out as described by Queiroz et. al. 2019, Trendel et. al. 2019 and Villanueva et. al. 2020 [32–34]. 200 μl of chloroform was added for each ml of TRIzol (1:5 v/v) to get biphasic separation. The mixture was vortexed and centrifuged for 15 min. at 12,000 × g at 4°C. The upper aqueous phase and the lower organic phase were carefully removed leaving the interphase in the tube. The interphase was again subjected to two extra phase separation cycles by adding 1 ml of TRIzol each time. After third cycle, the interphase was gently washed with RNase-free water and 0.1% SDS. The washed interphase was mixed with 4x SDS-PAGE sample buffer and proceeded for gel electrophoresis and Western blot analysis.

### CLIP-seq assay

UV cross-linking and interphase separation was done as described above. The interphase were solubilised with 1% SDS (RNase-free). Protein-bound RNAs of the solubilised interphase were then precipitated with isopropanol (1.2 volumes) and 3M sodium acetate pH 5.2 (1/10 volume). The pellet was washed with 100% ethanol followed by 70% ethanol and left for 5 minutes at room temperature to dry. The dried pellet were then solubilised in nuclease-free PBS. Simultaneously, protein A/G beads (Pierce) were cross-linked with the anti-PfRAP291 and anti-PfRAP070 rabbit antibodies using DSS cross-linker (Thermo Scientific™) as per the manufacturer’s protocol. The cross-linked protein A/G beads were mixed with the solubilised protein cross-linked RNAs (in PBS). RNasin™ (Promega) 2 μl/ml with DTT to a final concentration of 1 mM were used to prevent RNA degradation. The mixture was incubated for 3-4 hours at RT while mixing on a nutator. The beads were then washed three times with 0.1% PBST followed by one PBS wash. The washed beads were resuspended in 600 μl of proteinase K buffer (Tris-Cl 50 mM, EDTA 10 mM, NaCl 150 mM and 1% SDS) containing 4 μl of Proteinase K and incubated at 50 C for 1 hr with intermittent mixing. The mixture was centrifuged at 13,000 rpm for 20 minutes at 4 C and the supernatant was subjected to RNA isolation by TRIzol. This purified RNA was further used to prepare cDNA using iSCRIPT™ cDNA synthesis kit (Bio-Rad) according to the manufacturer’s protocol. These cDNAs were subjected to adaptor ligation and sequencing using Illumina NGS platform. The sequencing data was then subjected to CLIP-seq analysis. Raw reads were quality checked and adaptor trimmed by using FastQC (v.0.11.9) and Trim-Galore (v.0.6.6) [35,36]. The cleaned and processed reads were further aligned with *Plasmodium falciparum* 3D7 genome and human genome using BWA (v.0.7.17-r1188) and HISAT2 (v.2.21) [37,38]. Peak calling analysis was carried out by using Samtools (v.1.10) and PEAKachu (v.0.2.0) [39,40]. MA plots were generated by using the PEAKachu results. The MEME suit was used for the identification of motifs. R package, ChIPSeeker (v.1.18.0), was used to generate pie charts [41].

### Blue-Native PAGE

*P. falciparum* merozoites were isolated as described by Hill et. al., 2014 [29] and lysate was prepared using RIPA buffer. Briefly, Sample loading dye was prepared as mentioned in Wittig et. al., 2009 [42] and added to the merozoite lysate. The sample was run on Blue native-PAGE (4% stacking – 8% resolving) for overnight at 30 Volts in a cold room. Western blotting was performed and membrane was probed with anti-PfRAP070, anti-PfRAP291, anti-PfMSP-1_65_, anti-PfMSP-3N or anti-PfRhopH3b rabbit antisera 1:1,000 dilution for 2 hours. Blot was subsequently incubated with secondary antibody at 1:50,000 dilution and bands were detected using chemiluminescence detection kit.

### Far western assay

Far western assay was carried out according to the protocol described earlier [43]. Briefly 1-5 μg of recombinant proteins; rPfRAP070, rPfRAP291, rPfMSP-1_65_, rPfMSP-3N, rPfRhopH3b and a recombinant *Plasmodium* ring exported protein rPfREX (used as negative control), were run on SDS-PAGE individually and transferred to a membrane. The proteins on the membranes were denatured and renatured as described in Wu et. al., 2007 [43]. These membranes were blocked with 5% skimmed milk and incubated with 2 μg/mL of purified interacting bait proteins, i.e., recombinant PfRAP070 or PfRAP291 in protein-binding buffer (100 mM NaCl, 20 mM Tris (pH 7.6), 0.5 mM EDTA, 10% glycerol, and 1 mM DTT) for 4 hours at RT. Membranes were washed to remove the non-specific interactions and were incubated with rabbit anti-PfMSP-1_6_5 R2(1:1,000) or rabbit anti-PfMSP-3N (1:1,000) or rabbit anti-PfRhopH3b antisera (1:500) overnight at 4 °C followed by incubation with goat anti-rabbit HRP conjugated (1:30,000) or goat anti-mouse HRP conjugated (1:15,000) for 1 hour at RT. Finally, the blots were developed using ECL kit (Bio-Rad) and imaged with Chemi-Doc (Bio-Rad).

### Co-Immunoprecipitation assay

Briefly 10 μg of rPfRAP291 and rPfRAP070 were incubated with 10 μg of rPfMSP-1_65_, rPfMSP-3N or rPfRhopH3b proteins in separate reaction mixtures for 2 hours in 100 μl binding buffer (50 mM phosphate buffer at pH 7.0, 75 mM NaCl, 2.5 mM EDTA pH 8.0, 5 mM MgCl2, 0.1% NP-40 and 10 mM DTT). The reaction mixture was further incubated for 2 hrs at 4° C with 20 μl of Pierce protein A/G plus agarose beads crosslinked with 20 μg rabbit of rabbit anti-PfRAP070 or anti-PfRAP291 antisera [28]. The beads were then centrifuged at 1000 × g for 5 mins, washed with 200 μl of binding buffer containing 400 mM NaCl and boiled for 5 mins in SDS PAGE sample buffer. Proteins were subsequently electrophoresed, immunoblotted and probed with either mice anti-rPfMSP-165, anti-rPfMSP-3N and anti-rPfRhopH3b antisera followed by goat anti-mice HRP conjugated secondary antibody (1:100,000 dilution). The blots were developed using ECL kits (Bio-Rad) and imaged with Chemi-Doc (Bio-Rad).

### Indirect Immunofluorescence assay (IFA)

Confocal laser scanning IFAs were performed with *P. falciparum* blood stages. Cell fixation, antibody incubation and imaging were performed by standard techniques as described earlier (13). Liquid IFA was also carried out on fixed parasites for protein localization and co-localization studies following a protocol described earlier [44,45]. Blocking of merozoite stage parasite was carried in PBS containing 10% FBS (Foetal bovine Serum) for 2 hours. Parasites were further incubated with polyclonal anti-PfRAP070 or anti-PfRAP291 sera (1:100 dilution) in PBS containing 1% FBS for 3 h. Anti-peptide PfRAP070 and anti-peptide PfRAP0291 antibodies were used at 1:50 dilution in PBS containing 1% FBS for overnight. For co-localization studies anti-PfMSP-1_6_5 (1:100), anti-PfMSP-3N (1:100) and anti-PfRhopH3 (1:100) sera were used along with anti-PfRAP291 (1:100) or anti-PfRAP070 (1:100) sera. Subsequently, secondary rabbit or mice antibodies coupled with fluorophores Alexa 594 (red) and Alexa 488 (green) were used at 1:300 dilutions in PBS containing 1% FBS for 1 h. The DAPI was used for staining parasite nuclei. Finally, Microscopic examination was performed using a A1 confocal microscope (Nikon). Images were analysed using Nikon NIS Elements v 4.0 software. IMARIS image was created using the software IMARIS v 4.0.

### Invasion inhibition assay

Invasion inhibition assay was performed similarly as described earlier [27]. In brief, anti-PfRAP291 and anti-PfRAP070 heat inactivated sera’s were added to highly synchronized late trophozoite stage culture with *1%* parasitaemia at final concentration of 5, 10 and 20% in the *in-vitro* growth inhibition assay. Anti-PfMSP-1Fu anti-sera and pre-immune rabbit anti-sera were used as positive and negative controls at concentrations of 20% serum. The cultures were incubated for 40 hours for schizont rupture and merozoite invasion. Parasitaemia was counted by FACS (Fluro-scenes activated cell sorter). Percentage inhibition was calculated relative to the controls. Bars indicate mean ±SEM of triplicate measurements.

### RBC Binding Assay

Erythrocyte-binding assays were performed as recently described by Chourasia et. al., 2020 [28]. In brief, 10 μg of both recombinant RAP proteins fragments were incubated with 100 μl of fresh packed RBC for 1 hr. The RBCs were separated from the supernatant by spinning through 600 μl of Dibutyl phthalate (Sigma) at 12000 × g for 30 s. Proteins bound to the erythrocytes were eluted by incubation with 20 μl of 1.5 M NaCl in PBS at room temperature for 5 mins. Eluted proteins were mixed with equal volumes of 2x non-reducing sample buffer. The eluates were analysed by immune-blotting with respective antibodies. Recombinant PfClag9c fragment was used as positive control.

### Seroprevalence analysis

ELISA analysis was performed to determine the sero-reactivity of the proteins; rPfRAP070 and rPfRAP0291 using sera from naturally infected malaria patients as described earlier [46]. Sera from 28 *P. falciparum* malaria patients in India and 28 *P. falciparum* malaria patients in Liberia were used, while sera from 28 Danish volunteers was used as a negative control. The Danish volunteers were used to determine the positive threshold. Briefly, 96-well polystyrene flat-bottom plates (Nunc-Maxisorp; Thermo Scientific) were coated with 500 ng of rPfRAP291 and rPfRAP070 protein in carbonate-bicarbonate buffer and incubated overnight at 4 °C. Next day, 96 well plates were 3 times washed with PBST (PBS containing 0.1% Tween buffer and blocked for 2 hrs in 5% milk. After a blocking step, the wells were incubated with Indian, Liberian, or Danish serum samples (1:200 dilution) at RT for 1 h, followed by incubation with HRP-conjugated goat anti-human IgG (1:3,000; Sigma) for 1 h at RT. The bound antibody was detected with tetramethylbenzidine (TMB) solution (Sigma). Plates were extensively washed between each incubation period with PBST (0.1% Tween 20).

### Statistical analysis

Graphs and statistical analysis were performed with Microsoft Excel and GraphPad Prism, version 7. Significance was calculated using Student t-test.

## Results

### Genome-wide screening for RAP domain containing proteins in *Plasmodium falciparum*

Using various bioinformatic tools, 21 ortholog groups having RAP domain have been identified in genomes of different *Plasmodium* species. The sequences of all the 20 RAP domains present in the *P. falciparum* RAP domain containing proteins were extracted from PlasmoDB and UniProt databases. These sequences were aligned using T-COFFEE software version-11 (with default settings) as represented in supplementary figure S1-A. Through MEME analysis, a 28 amino-acids long common motif (represented in red box) LxxxGxxxxxxW was identified with **L**(leucine), **G**(glycine), and **W**(tryptophan) as highly conserved residues (Fig. S1-A). The p-value for the common motif identified in respective proteins is mentioned in supplementary figure S1-B. The size of the RAP domain in different *P. falciparum* proteins ranged from 56 to 73 amino acids with the majority having approximately 60 amino acid residues. Interestingly, the RAP domain was always positioned at the C-terminal end of these proteins.

To understand the role(s) of RAP domain proteins in *P. falciparum*, we decided to characterize two RAP domain proteins; PfRAP291 and PfRAP070. Sequences similar to PfRAP291 and PfRAP070 (about 90% and 64% similarity respectively) were identified in other *Plasmodium* species and in other apicomplexans, but not in primates. RAP domain is present at C-terminal of these proteins as observed for other *P. falciparum* RAP domain proteins (Fig. S1-C & S1-D in the supplemental material).

### Expression and purification of recombinant PfRAP291 and PfRAP070 protein fragments in *Escherichia coli*

The rPfRAP291_876-1311_ and rPfRAP070_2088-2925_ gene fragments were expressed in *Escherichia coli* Shuffle 30 cells and were analysed by SDS-PAGE followed by western blot analysis using anti-His antibody (Fig. 1A-i, and Fig. S2-A & Fig. S2-B). Sub-cellular fractionation studies revealed the presence of both recombinant proteins in soluble and as well as in cell pellets. Recombinant PfRAP291 and PfRAP070 proteins were purified from the soluble fraction to near homogeneity on a Ni-NTA^+^ column under non-denaturing conditions (Fig. 1A-ii). In case of rPfRAP291 protein, two bands of sizes; ~28 kDa and ~70 kDa were observed, while purified rPfRAP070 protein showed a single band of ~29 kDa. LC-MS/MS analysis of these bands confirmed the expression of rPfRAP291 and rPfRAP070 protein (Figure S-2C in supplemental material). Antibodies to these two RAP domain protein fragments; rPfRAP291 & rPfRAP070 were generated in mice and rabbits. We also produced anti-peptide antibodies to these proteins as indicated in figure. 1A (i).

**Fig. 1.**
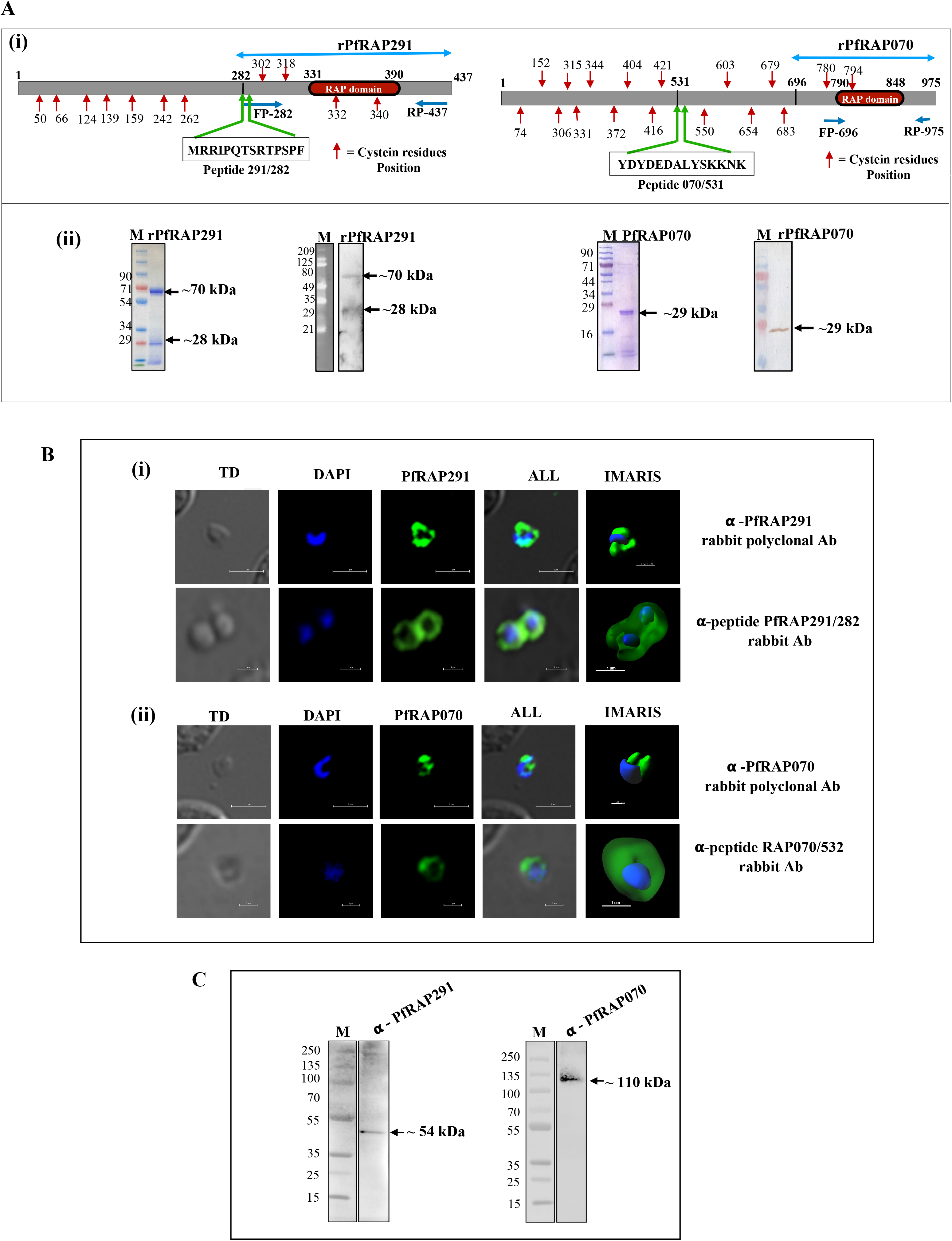
Expression of recombinant proteins, rPfRAP291 and rPfRAP070, and the expression analysis of native PfRAP291 and PfRAP070 at *P. falciparum* asexual blood stages. **(A-i).** Schematic showing the organization of PfRAP291 and PfRAP070, two *P. falciparum* RAP domain proteins. Double-head blue arrows indicate the size of protein fragments expressed. Boxed sequences show the peptide sequence used to generate peptide antibodies for each protein. FP-forward primer; RP-reverse primer (**A-ii).** Coomassie stained SDS-PAGE and western blot analysis showing the purified rPfRAP291 and rPfRAP070 protein fragments. Western blot analysis were performed using anti-His antibody. (**B)** Immunofluorescence assays (IFA) followed by confocal microscopy showing localization of PfRAP291 **(B-i)** and PfRAP070 **(B-ii)** proteins at *P. falciparum* merozoite stage. Scale Bar represents 1 μm. (**C)** Western-blot analysis showing the expression of native PfRAP291 **(C-i)** and PfRAP070 **(C-ii)** proteins at *P. falciparum* asexual blood stage (merozoite).

### PfRAP291 and PfRAP070 proteins are expressed at asexual blood stages

To study the expression and localization of PfRAP291 & PfRAP070 proteins at asexual blood stages, immunofluorescence staining of fixed *P. falciparum* parasites and western blot analysis for parasite lysate were performed using anti-PfRAP291 and anti-PfRAP070 antibodies. As shown in figures 1B & supplementary figures S3-A & S3-B, a strong immunofluorescence staining was observed at merozoite, trophozoite and schizont stages for each of the two RAP domain proteins. We could observe typical surface staining for each of these proteins similar to the one reported for merozoite surface proteins [27,47]. In addition to surface staining, staining was also seen in parasite cytoplasm at trophozoite and schizont stages. To take care of specificity of staining, we also raised antibodies to peptides corresponding to each proteins. Similar staining pattern was observed using anti-peptide specific anti-bodies in merozoites (Fig. 1B). To further study the expression of these RAP proteins at merozoite stage, we next performed western blot analysis using merozoite lysate with anti-PfRAP291 or anti-PfRAP070 rabbit antibodies. An immune-reactive band corresponding to the expected size of each of the two native proteins was seen in parasite lysates of merozoites (Fig. 1C). None of these bands were observed with pre-immune sera (see Fig. S3-A-(ii) and S3-B-(ii) in the supplemental material). We next performed the LC-MS/MS analysis of the native bands by extracting proteins from gel pieces. As shown in figure S3-C, we were able to identify the native proteins in these gel bands. Together, these results demonstrate that PfRAP291 & PfRAP070 proteins are expressed at merozoite stage of the parasite.

### PfRAP291 and PfRAP070 bind *P. falciparum* RNA(s)

Although the name “RAP” domain stands for “RNA binding domain abundant in Apicomplexan”, there has been limited work to show the RNA species that bind to these RAP domain proteins [17,18]. To know whether the PfRAP291 or PfRAP070 proteins binds RNA(s), parasites were irradiated with 254 nm UV light and cell pellets were analysed on SDS-PAGE followed by western blot analysis using anti-PfRAP291 and anti-PfRAP070 antibodies (Fig. 2A). As shown in figures 2B & 2C, after UV cross-linking step, we could observe both the proteins moving higher than their migration in un-crosslinked lanes. These results were further confirmed when we performed an orthogonal organic phase separation method (OOPS) on UV cross-linked lysate of *P. falciparum* late schizont stage to study the RNA-protein interactome [32–34]. The interphase was collected and protein-RNA complexes were further enriched by two subsequent TRIzol steps and was analysed for the presence of two *P. falciparum* RAP domain containing proteins; PfRAP291 and PfRAP070. We failed to get sufficient amount of interphase in non-cross linked lysate. Protein-RNA conjugates present in the enriched interphase were analysed by western blotting using specific antibodies. Schematic of the protein-RNA separation is presented in figure 2A. As shown in figure 2D, we could enrich these RNA cross-linked proteins in the interphase layer that were detected in western blot analysis, thereby suggesting interactions of the two RAP domain proteins with RNAs. This was specific as PfBiP which was used as a negative control could not be identified in the interphase (Fig. S4). Together, these results indicated that both PfRAP291 and PfRAP070 bind some RNA species of *P. falciparum* origin.

**Fig. 2.**
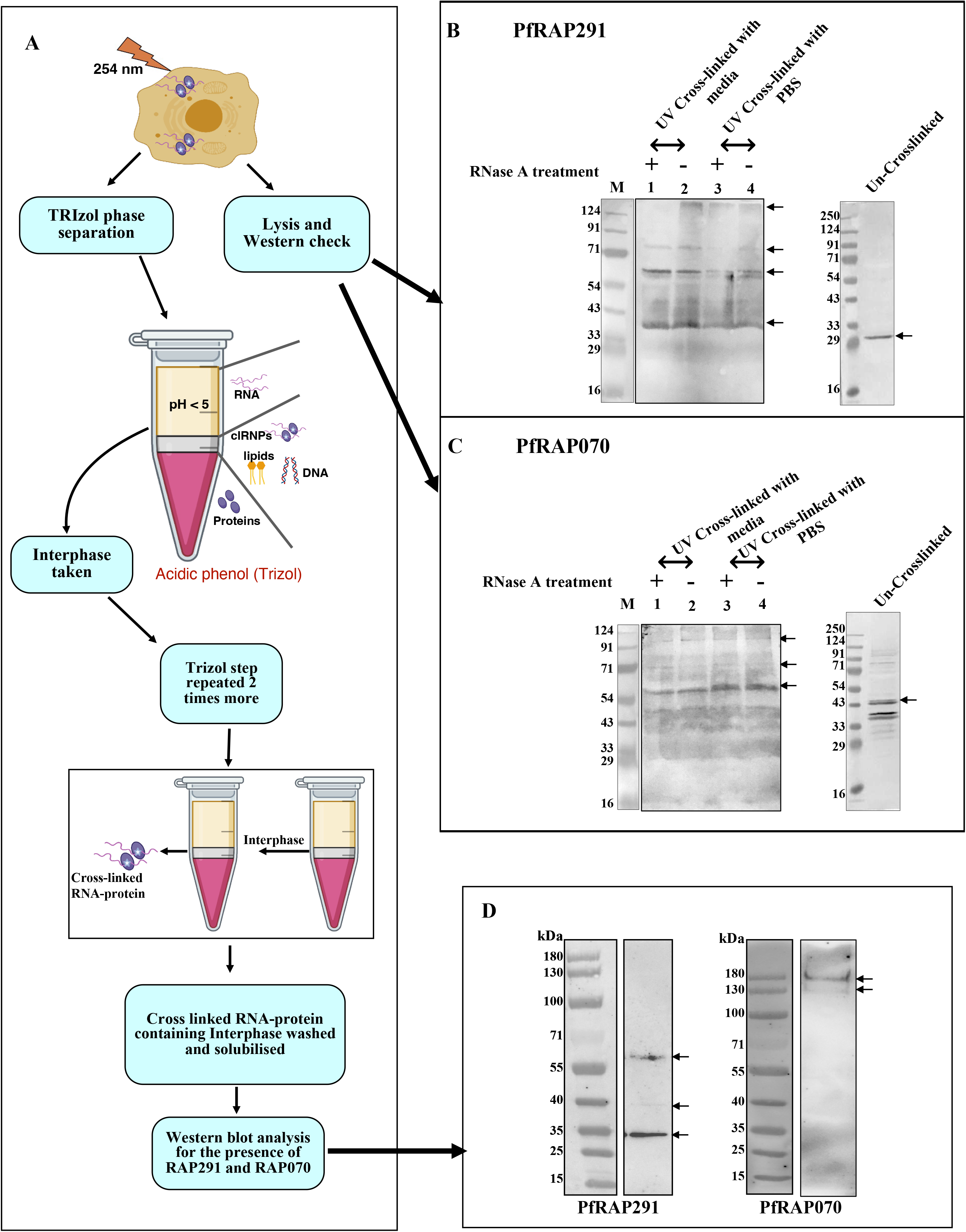
Identification of RNA-protein interactions using orthogonal organic phase separation. **(A)** Schematic representation of the UV cross-linking and organic phase separation procedures. **(B & C)** Western-blot analysis showing electromobility-shift of RAP domain proteins **(B**:PfRAP291 & **C**:PfRAP070) on SDS-PAGE after UV cross-linking. **(D)** Western-blot analysis showing the presence & enrichment of PfRAP291 and PfRAP070 proteins in the purified TRIzol interphase after UV cross-linking of the parasites.

### Identification of RNAs bound to RAP proteins

To identify RNAs interacting with PfRAP291 and PfRAP070, we performed CLIP-seq experiment in *P. falciparum*. Briefly, protein-RNA conjugates were isolated from interphase as described earlier. These protein-RNA complexes were immunoprecipitated using anti-PfRAP291 or anti-PfRAP070 rabbit antibodies. RNAs were released from immuno-precipitates by proteinase K treatment and were used to prepare cDNAs. These cDNAs were then subjected to adaptor ligation and sequencing using the high-throughput sequencing Illumina NGS platform. CLIP-seq experiments were performed in duplicates. Schematics of isolation of protein-RNA complex and CLIP-seq is presented in figure 3A. The MA plot, generated using DESeq2 after initial peak calling (using PEAKachu), shows the general trend of the log2 fold-change in dependence of the average mean of the expression rate of the peaks where red dots are significant peaks (Fig. 3B-i & ii). Approximately, 83% and 52% of the reads from PfRAP070 and PfRAP291 immunoprecipitates, respectively, were mapped to *P. falciparum* genome (Additional file1 in supplementary materials). A very low read alignment rate of ~4-8% was obtained in the human genome (Figure 3B). In total 43,139,186 (for PfRAP291) 50,285,370 (for PfRAP070) peaks sequences were detected and of these peak sequences 22,406,880 (for PfRAP291) and 41,908,250 (for PfRAP070) were found as unique peak sequences in *Plasmodium* (Fig. 3B-iii). Reproducible peaks from RNA seq data were annotated with their location in different *P. falciparum* genomic regions (5’ untranslated, 3’ untranslated, coding & intragenic sequences (Fig. 3C and Additional File-1). Among the top hits are the sequences that are part of 28s and 18s ribosomal RNAs (Tables 1&2 and Additional File-1). Based on the CLIP-seq analysis, several consensus motifs were identified for potential binding to these RAP proteins. The sequences of top enriched peaks from each of the two samples were used to search for enriched motifs. Figure 3D shows the top 5 motifs identified as recognition element for PfRAP291 & PfRAP070 proteins. Top twenty peak sequences identified in *P. falciparum* genome for each of the two RAP proteins CLIP-seq analysis showed that these RNAs are mainly associated with 28s (large subunit) and 18s (small subunit) ribosomal RNA (Table.1 & Table.2). Together, these results point towards a role of these proteins in ribosome assembly and/or functions.

**Fig. 3.**
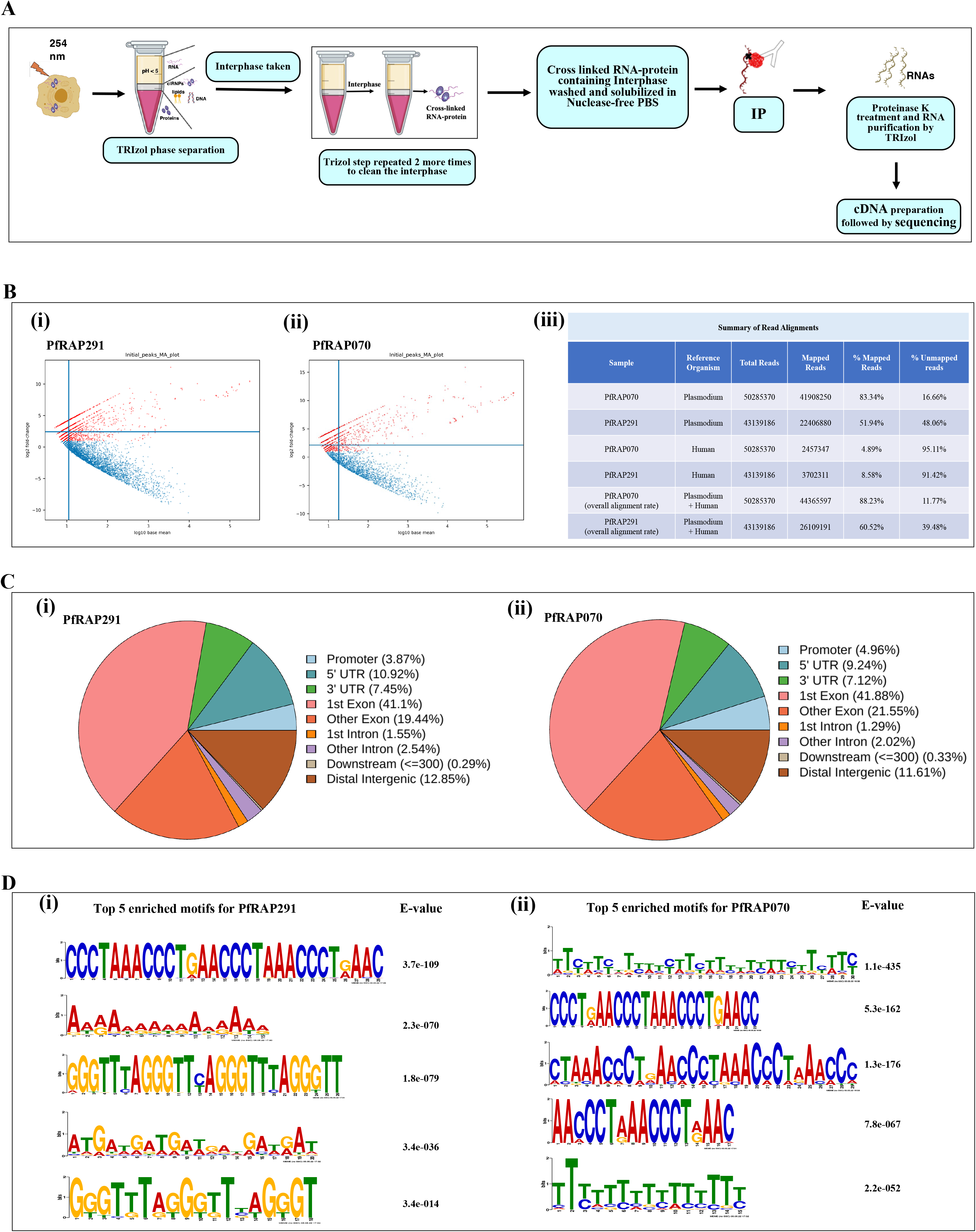
CLIPseq analysis reveals binding of RNAs with PfRAP291 and PfRAP070. **(A)** Schematic illustrating the key steps of CLIPseq protocol. **(B)** MA plots (i and ii) of initial peaks called using PEAKachu for *Plasmodium falciparum*. MA stands for M (log ratio) and A (mean average). The blue lines depict the normalisation constants. Red dots are significant peaks. The table (iii) shows the summary of all read alignments. **(C)** Representative chart illustrating the proportion of peak distribution in 3’UTR, 5’UTR, exons, introns, downstream regions and distal intergenic regions for PfRAP291(i) and PfRAP070(ii) bound RNAs. **(D)** Top 5 consensus motifs identified by CLIPseq analysis for PfRAP291(i) and PfRAP070(ii).

**Table 1.**
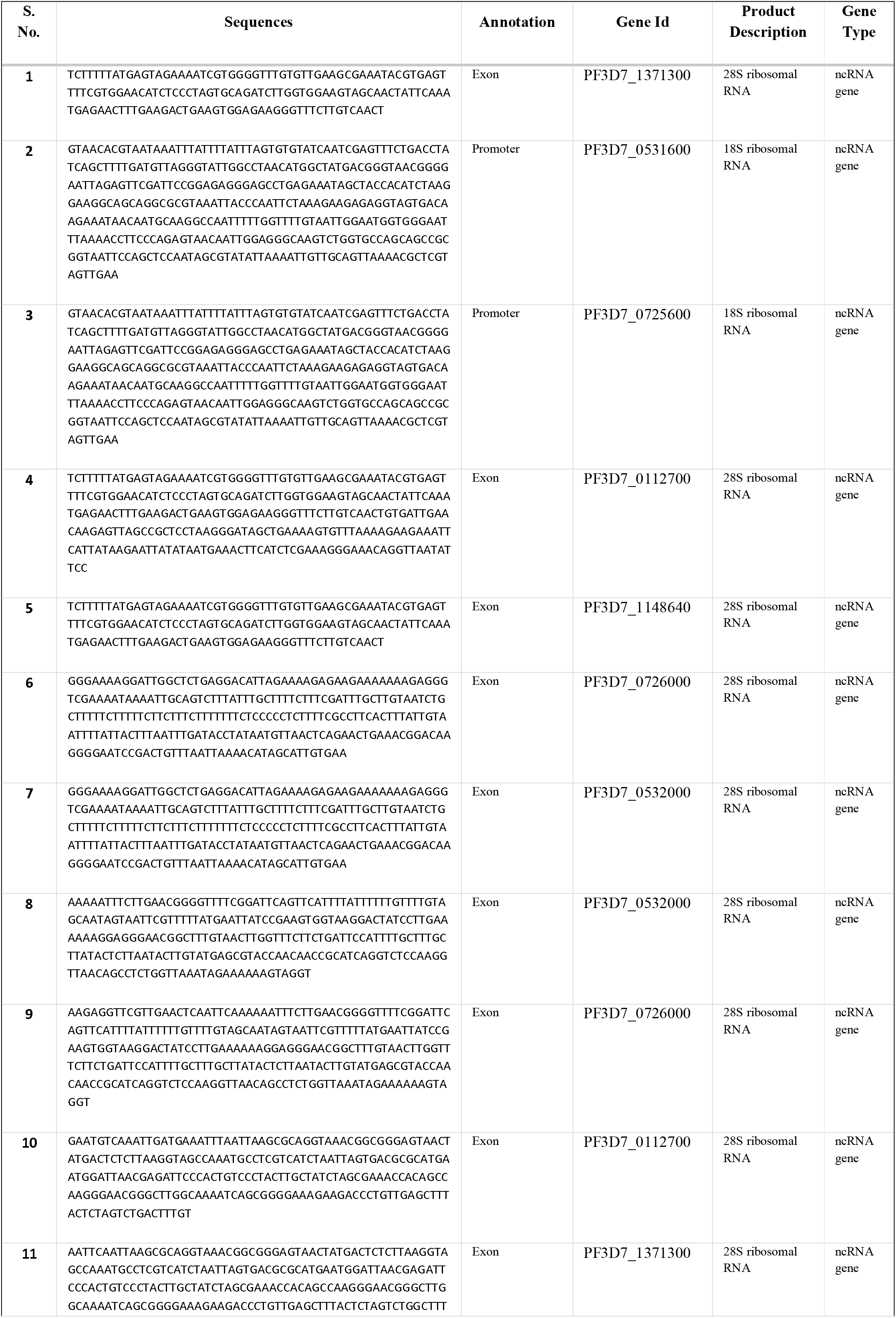
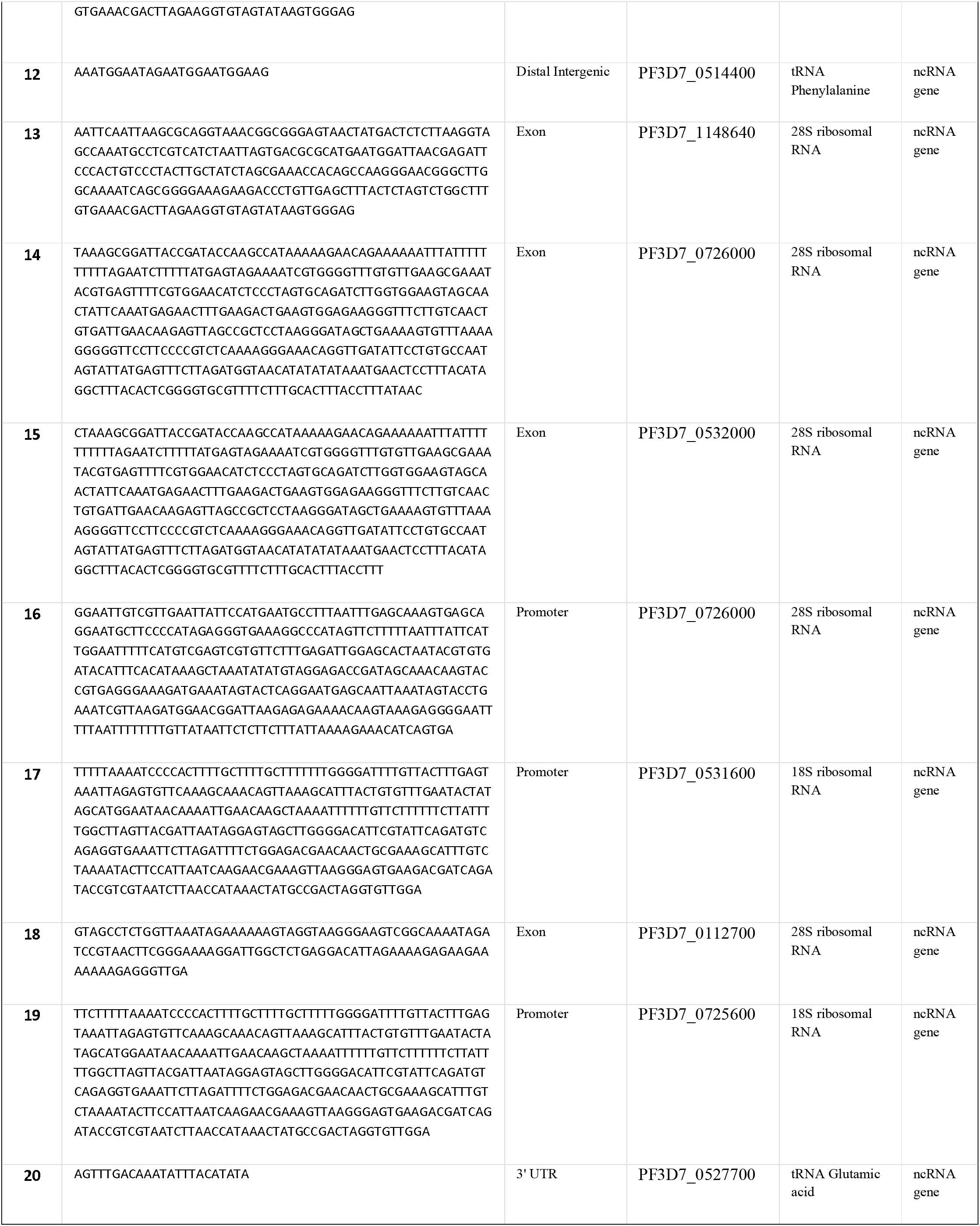
Top 20 peak sequences for PfRAP291 after CLIP-seq analysis.

**Table 2.**
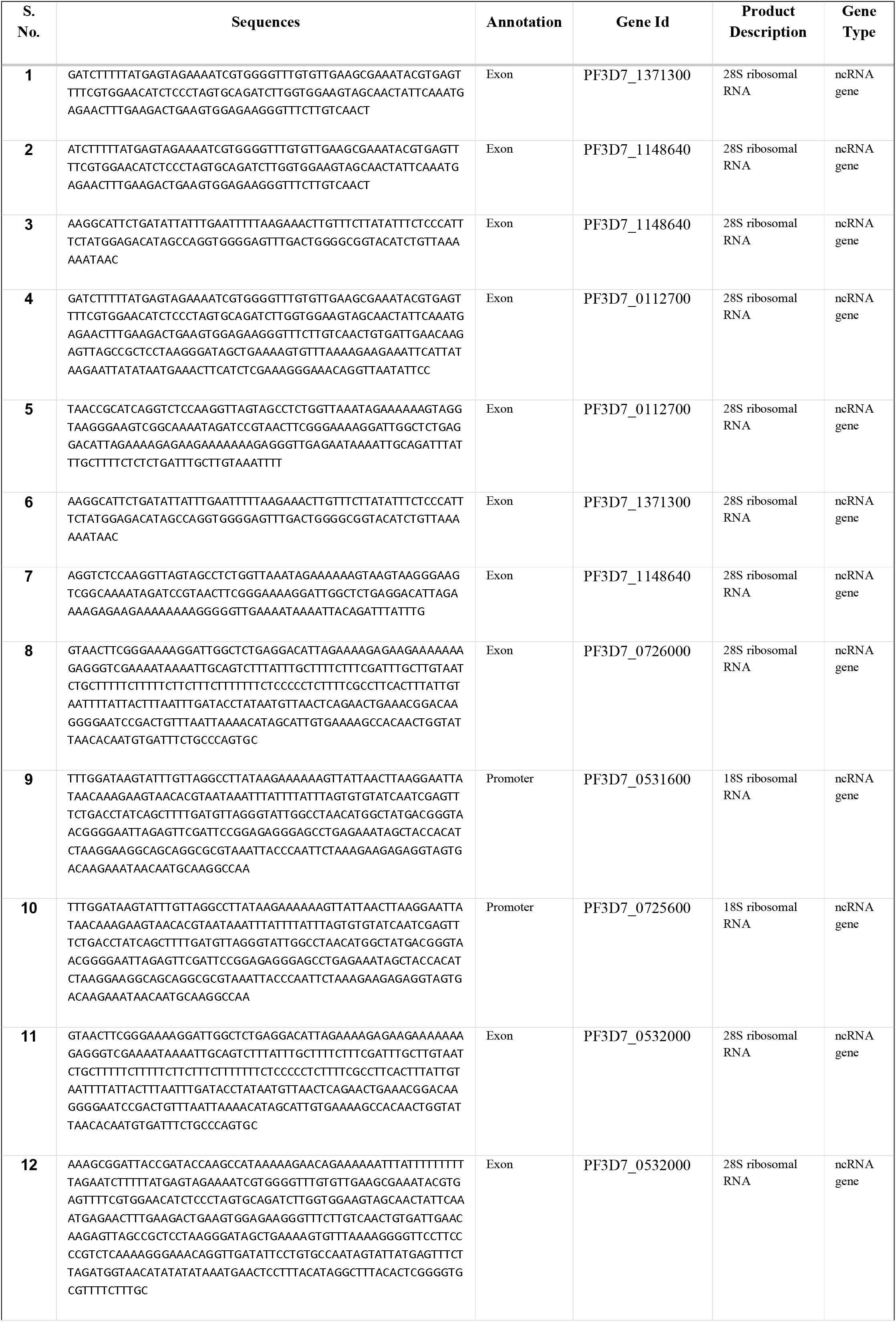
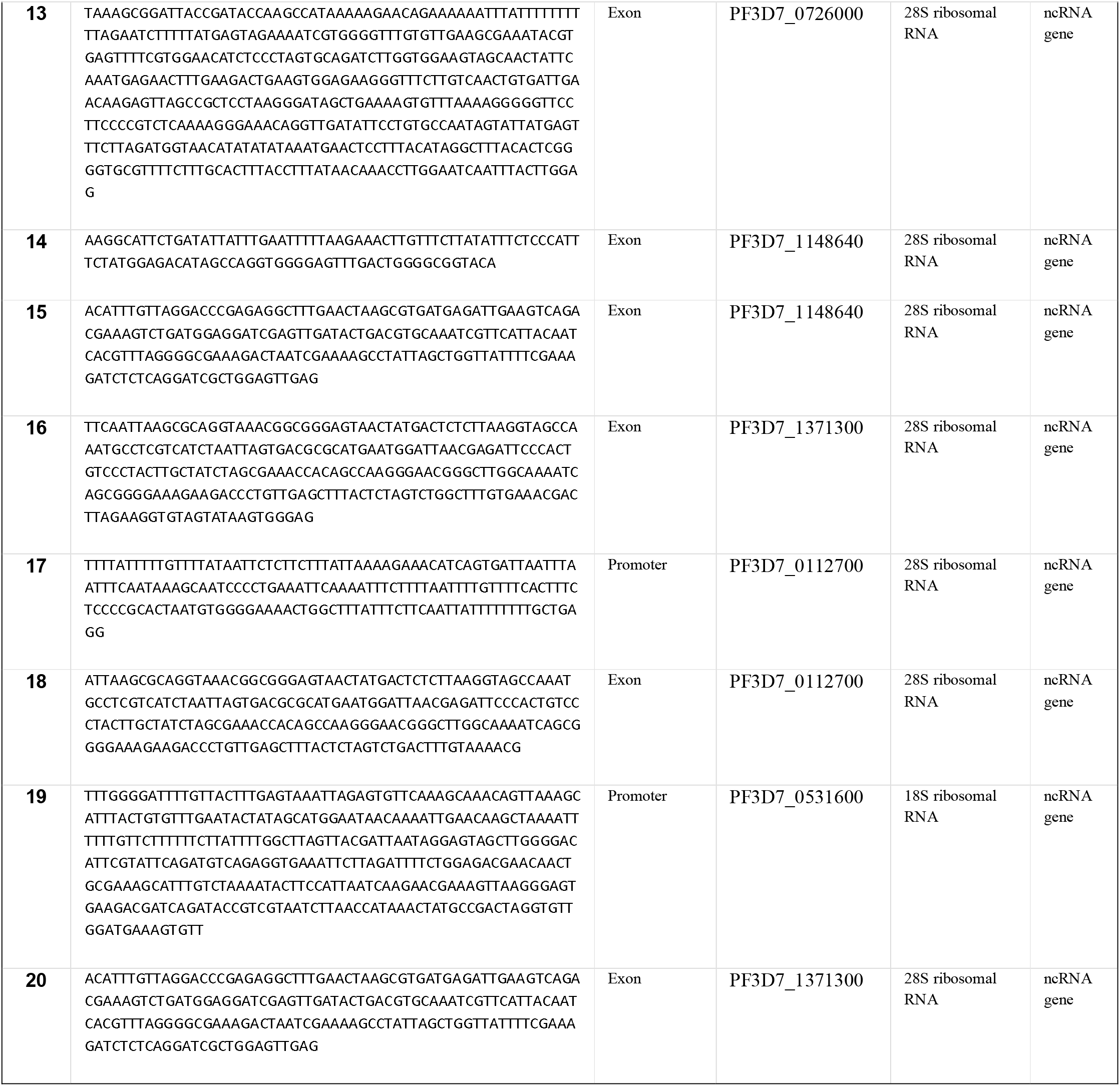
Top 20 peak sequences for PfRAP070 after CLIP-seq analysis.

### Association of PfRAP291 & PfRAP070 with MSP-1 complex

Since these two RAP protein are expressed on merozoite surface, we next characterized these proteins for their association with other merozoite surface proteins. A Blue Native PAGE analysis of merozoite membrane extract followed by western blot analysis using anti-PfMSP-1_65_, anti-PfMSP-3N, anti-PfRhopH3b, anti-PfRAP291 or anti-PfRAP070 antibodies was performed. As shown in figure 4A, both RAP proteins existed in two complexes of molecular sizes ~650 kDa and ~800 kDa along with MSP-1, MSP-3 and RhopH3 proteins. To study the interaction among the components of these complexes, we next performed Far-western blot and co-immunoprecipitation analysis using a protocol described earlier [43]. For these interactome analysis, three recombinant proteins; rPfMSP-165, rPfMSP-3N and rPfPfRhopH3b were generated as described earlier (see supplementary Fig. S5A) [27,28,47]. For the Far-western blot analysis either of the two recombinant RAP proteins, which served as a bait proteins were run on a SDS-PAGE and blots were probed with each of the three recombinant prey proteins; rPfMSP-1_65_, rPfMSP-3N or rPfRhopH3b. After the incubation for 1 h, each blot was developed using respective antibodies. As shown in figure S5B, all the three recombinant proteins; rPfMSP-1_65_, rPfMSP-3N and rPfRhopH3b interacted well with each of the two RAP proteins, thus confirming interactions among these proteins. These interactions were specific as a non-related recombinant protein, rPfREX (ring exported protein-1), which is not part of the MSP-1 complex, did not show interactions with any of the two RAP domain proteins. For the *in vitro* co-immunoprecipitation analysis, rPfRAP291 and rPfRAP070 proteins were incubated with each of the three recombinant bait proteins; rPfMSP-1_65_, rPfMSP-3N and rPfRhopH3b separately in a binding buffer as described in materials and methods. Bound proteins, if any, was immunoprecipitated using either anti-PfRAP291 or anti-PfRAP070 antibodies. Immunoprecipitates were run on SDS-PAGE and western blotting was performed using either anti-MSP-165 or anti-MSP-3N or anti-RhopH3b antibodies. As shown in figure 4B, each of the three MSP-1 complex associated proteins were recognized by their respective antibodies in the immuno-precipitates, confirming the association of the two RAP proteins with the components of the MSP-1 complex. Recombinant PfREX protein served as a negative control and it did not show interaction with either of the two RAP proteins (Fig. 4B)

**Fig. 4.**
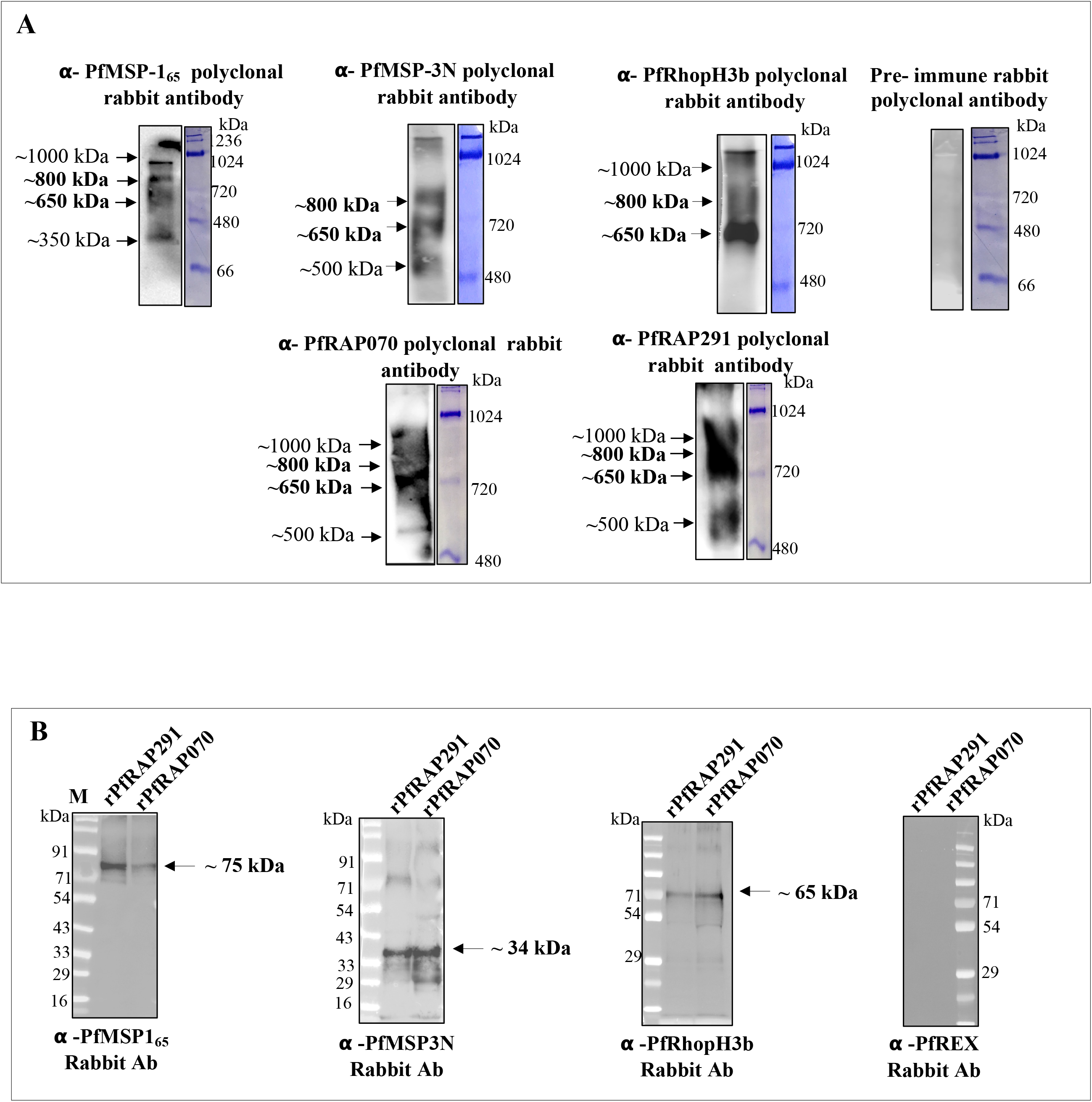
PfRAP291 and PfRAP070 are associated with a MSP-1 associated complex. **(A)** Blue Native PAGE followed by western blot analysis of merozoite lysate, using anti-MSP-165, anti-MSP-3N, anti-RhopH3b, anti-PfRAP291 and anti-PfRAP070 rabbit sera. All the sera recognized common ~650 kDa and ~800 kDa bands, while pre-immune sera failed to recognize any such band. **(B)** Co-immunoprecipitation analysis showing the interaction of the two RAP domain proteins, rPfRAP291 and rPfRAP070, with the proteins of MSP-1 complex; rPfMSP-165, rPfMSP-3N and rRhopH3b.

To further illustrate the association of PfRAP291 & PfRAP070 proteins with MSP-1 complex, co-localization studies for these proteins were performed on fixed merozoites/schizonts in liquid cultures by immunofluorescence staining using their specific antibodies. PfRAP291 partially co-localized with MSP-1, MSP-3 and PfRhopH3 proteins with a Pearson’s coefficients of more than 0.6 at merozoite and schizont stages, advocating the co-existence of PfRAP0291 with proteins of MSP-1 complex on the merozoite surface (Fig. 5A and Fig. S6-A). Like-wise PfRAP070 co-localized with MSP-1, MSP-3 as well as with PfRhopH3 with a Pearson’s coefficients of more than 0.6 at merozoite and schizont stages (Fig. 5B and Fig S6-B). PfRAP291 and PfRAP070 also co-localized well with each other as well (see Fig. S7-A & S7-B in the supplemental material). Together, these results unequivocally showed the presence and association of the two RAP domain proteins, PfRAP291 and PfRAP070, on the merozoite surface in a large MSP-1 associated complex.

**Fig. 5.**
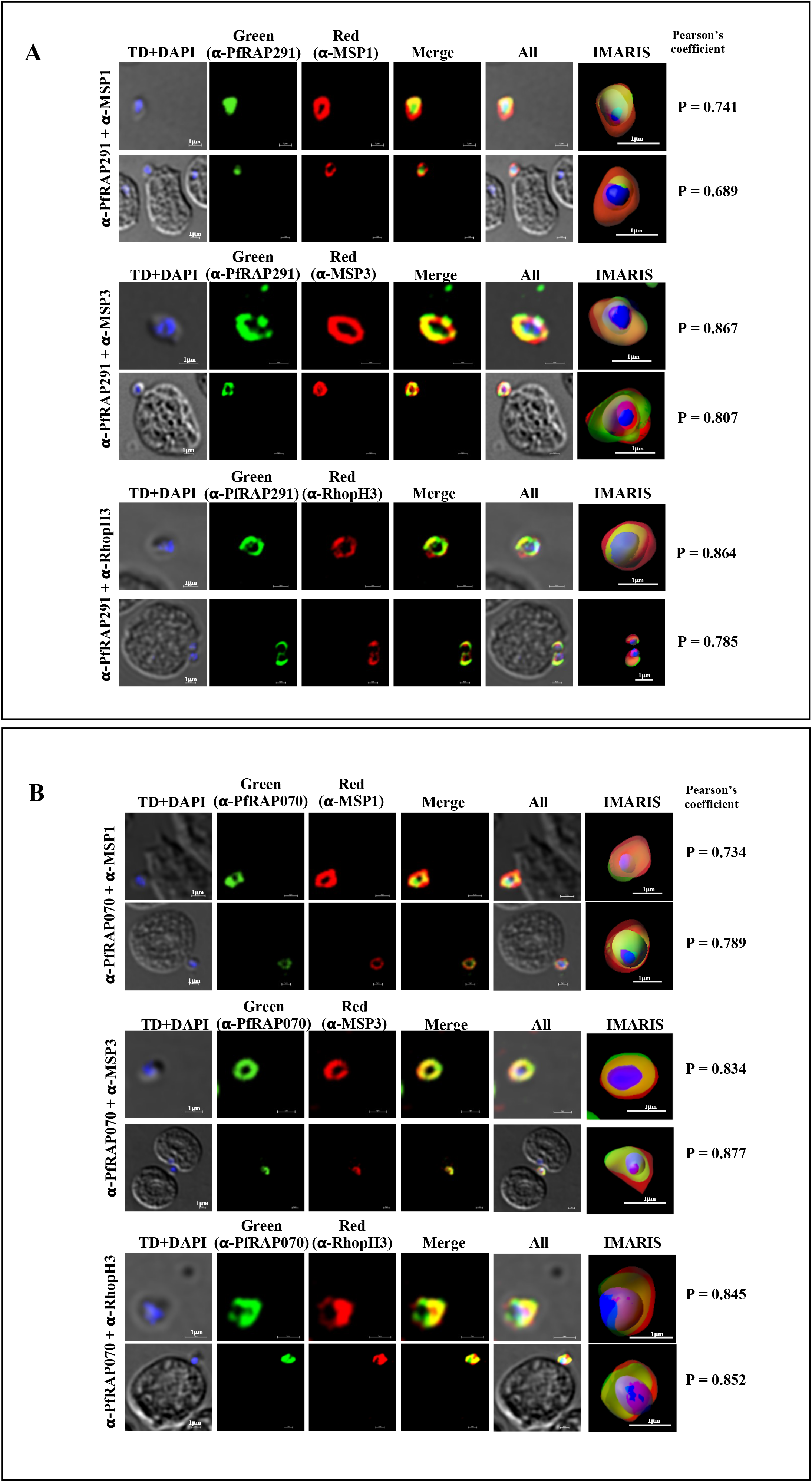
Co-localization studies on *P. falciparum* merozoite stage parasites using immunofluorescence assay: Immunofluorescence assays (IFA) followed by confocal microscopy imaging. **(A)** IFA images showing co-localization of PfRAP291 with PfMSP-1, PfMSP-3 and PfRhopH3. **(B)** IFA images showing co-localization of PfRAP070 with MSP1, MSP-3 and PfRhopH3b. P = Pearson’s coefficient; TD = transmitted light channel. IMARIS software was used to convert confocal images into clear informative schematics. Scale bar represents 1 μm.

### Humoral immune responses to the RAP proteins associated with MSP-1 complex and invasion inhibition assay

To know whether PfRAP291 or PfRAP070 are immunogenic during natural infections, seroprevalence analysis was performed by ELISA for these proteins using plasma from Africa and India. GLURP-R_o_ protein [26] was used as a positive control. Recombinant PfRAP291 & PfRAP070 proteins were frequently recognized by sera from Liberia with seropositivity rates of 78.6% and 49% respectively. Interestingly, these antigens were also recognized by sera from India with seropositivity rates of 32.10% & 17.90% respectively (Fig. 6A). The lower seropositivity rates observed among Indian samples may be related to a lower transmission intensity in this area compared with that of Liberia. Indeed, the GLURP-R_O_ seropositivity rate was 92.8% & 32.10% in Liberian and Indian sera respectively (Fig. 6A).

**Fig. 6.**
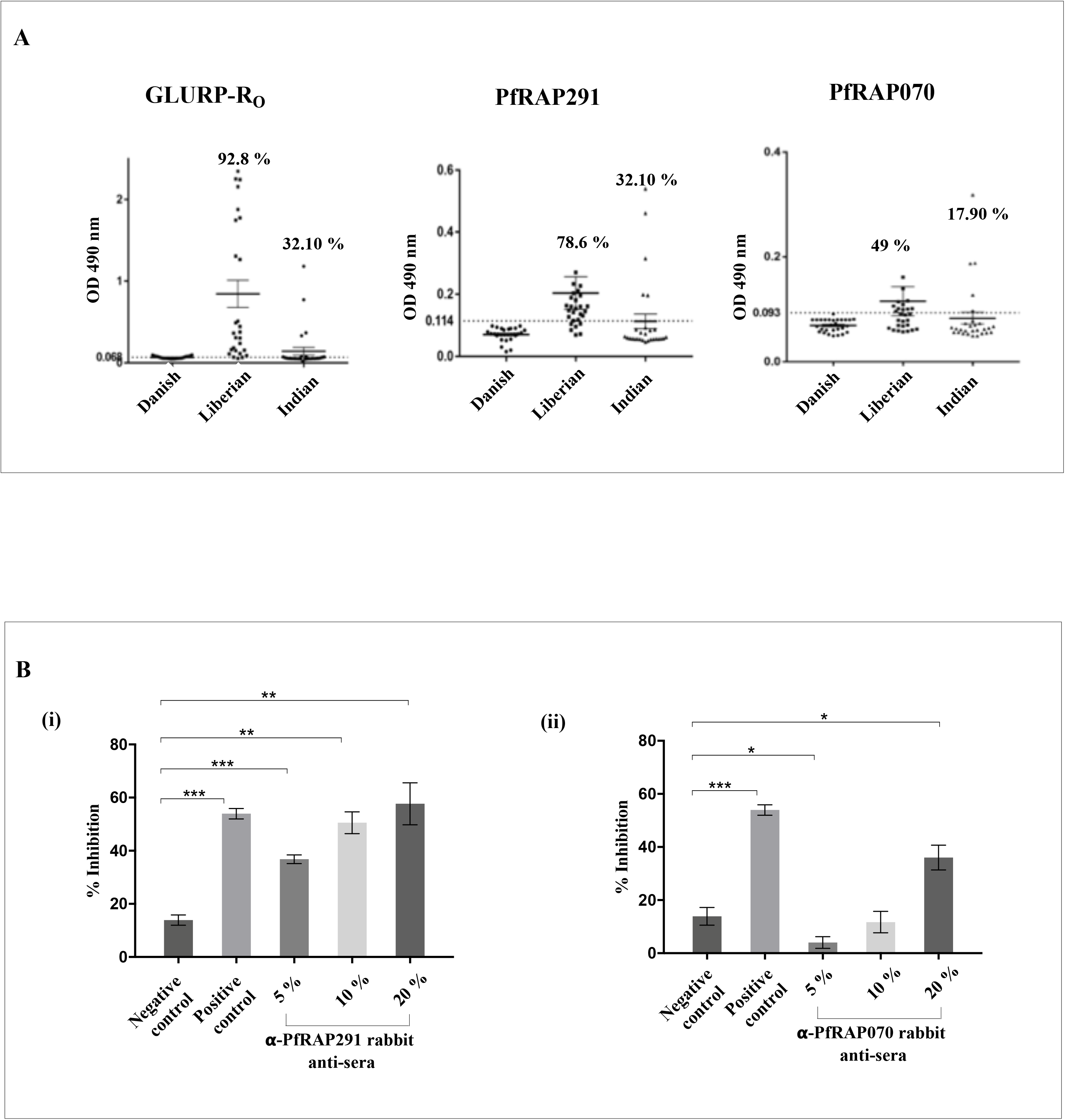
Naturally acquired immune responses and vaccine potential of PfRAP291 and PfRAP070 proteins. **(A)** Naturally acquired humoral antibodies against rPfRAP291 and RAP070 proteins in Indian and Liberian populations. ELISA was performed to analyse seroprevalence against RAP proteins in 28 naturally infected sera from areas of malaria endemicity in Liberia and India. The dotted line represents positivity threshold limits calculated by mean relativities twice the SEM results from 28 serum samples from Danish nonimmune sera. **(B)** Bar graph representing invasion inhibition using **(i)** anti-PfRAP291 and **(ii)** anti-PfRAP070 rabbit sera in concentration dependant manner. FACS analysis was done to determine parasite after 40 h of incubation. Data represents the means of 3 replicates, and error bars represent standard errors of the means. Significant results are indicated as follows: ***, p≤ 0.005; **, p≤ 0.05; *, p≤ 0.05. All other comparisons show no significant differences.

We next evaluated the inhibitory potential of anti-PfRAP291 or anti-PfRAP070 sera on the invasion of RBCs by *Plasmodium* merozoites at different sera concentrations. Rabbit anti-sera against the two RAP domain containing proteins were added at concentrations of 5%, 10% and 20% in tightly synchronized *P. falciparum* culture at the late trophozoite stage. At 40 h post-infection (hpi), parasites were Ethidium bromide (Etbr) stained and analysis was carried out by fluorescence activated cell sorter (FACS). Anti-PfRAP291 specific rabbit anti-sera moderately inhibited parasite invasion with 36.8%, 42.49% and 47.97% efficacy respectively (Fig. 6 B-i). Anti-PfRAP070 rabbit sera showed inhibitory potential of 36.04% at 20% serum concentration (Fig. 6 B-ii). Anti-MSP-1_Fu_ IgG [48] served as positive control and showed an inhibition of ~55.33% at 20% serum concentration. Pre-immune rabbit sera was taken as negative control which showed 13.5% inhibition at 20% serum concentration. Together, these results showed that anti-rPfRAP291 or anti-rPfRAP070 antibodies moderately inhibit the parasite invasion.

### rPfRAP291 or rPfRAP070 protein fragments do not bind human RBCs

To know whether any of the two RAP protein fragments generated in this study, binds to human RBCs, an *in vitro* erythrocyte binding assay was performed. PfClag9c protein fragment that has been previously shown to bind human RBCs served as a positive control [28]. Briefly, three recombinant proteins were incubated with washed human erythrocytes in an independent reaction mixture and bound proteins were analysed by western blot using specific antibodies. As shown in supplementary figure S8, neither PfRAP291 or PfRAP070 showed specific interaction with erythrocytes in comparison to PfClag9c that showed nice specific binding to human erythrocytes.

## Discussion

Apicomplexan parasites including *Plasmodium* spp. express several RAP domain proteins and only few of these proteins have been characterized. Here, we studied two RAP domain containing proteins; PfRAP291 and PfRAP070 for their expression and their binding to parasitic RNAs at asexual blood stages of *P. falciparum*. To get insights into the role of PfRAP291 or PfRAP070 at asexual blood stage, particularly in the invasion process, a segment of each of these proteins was expressed in *E. coli*, proteins were purified to > 90% homogeneity and specific antibodies were raised against these proteins. These antibodies recognized specific bands in asexual blood stage of *P. falciparum* lysates and stained the merozoite surface. Pattern of staining on *P. falciparum* schizont stage was typical of a merozoite surface protein [27,49–51]. In addition we also generated antibodies specific to peptides corresponding to these proteins and similar staining pattern was seen with anti-peptide antibodies.

Since these two proteins possess “RAP domain”, a putative RNA binding domain, we next looked for their interaction with parasite RNA(s). Studies in mammalian cells and *Chlamydomonas reinhardtii* have confirmed the role of RAP domain containing proteins in RNA metabolism [17]. For example, FASTK (Fas activated serine/threonine kinase), a RAP domain containing protein, has been demonstrated to play an essential role in regulation of mitochondrial ND6 mRNA levels [20,21]. Like-wise knock-out of FASTK4 has also suggested its involvement in mitochondrial RNA processing [21]. Recently using eCLIP-seq methodology in *Plasmodium*, two RAP proteins; PfRAP01 (PF3D7_0105200) and PfRAP21 (PF3D7_1470600) have been shown to bind distinct mitochondrial rRNA transcripts associated with the small and large mitoribosome subunits [22]. To provide evidence for PfRAP291 or PfRAP070 protein binding to RNAs, if any, we performed RNA-protein interaction assay in *P. falciparum* parasites using orthogonal organic-phase separation (OOPS) method [32–34], as well as CLIP-seq analysis of specific immunoprecipitates of crosslinked parasite lysates after phase separation [34]. True to the presence of RAP domain with predicted RNA binding property, we were able to identify RNA-RBPs (RNA-RNA binding proteins) interactions in *P. falciparum* parasites for both the RAP proteins, thus confirming the interactions of PfRAP291 and PfRAP070 proteins with parasitic RNA(s). Analysis of CLIP-seq data revealed that many of these RAP proteins binding RNAs can be mapped to 28s & 18s ribosomal RNAs associated with large and small ribosome subunits, thus suggesting the role(s) of these proteins in ribosome functions and/or assembly. These results, in concordance with a previous report by Hollin et. al., suggest that *Plasmodium* RAP proteins are playing important role in regulation of ribosome and/or mitoribosome assembly/functions [22].

Based on localization of these RAP domain proteins in parasite cytoplasm and on merozoite surface, we next looked for their association with parasite merozoite surface complex(s). A blue native page analysis of *P. falciparum* schizont stage parasite lysate followed by western blot analysis using anti-PfRAP291, and anti-PfRAP070 antibodies along with antibodies against three components of MSP-1 complex; anti-PfMSP-1, anti-PfMSP-3 and anti-PfRhopH3 antibodies revealed that these RAP proteins are part of ~650 kDa and ~800kDa MSP-1 complexes. These results were further corroborated by Far-Western and co-immunoprecipitation analysis. Together these results suggest that the two RAP proteins studied here are multifunctional proteins; having a role in ribosome regulation and also may have a role in merozoite invasion of RBCs or in parasite survival.

As these two RAP proteins were localized on merozoite surface, we further analysed the seroprevalence of their antibodies in *P. falciparum* infected individuals and also tested the ability of anti-PfRAP291 and anti-PfRAP070 antibodies in parasite invasion inhibition assays. Low to moderate seroprevalence was observed for both the RAP proteins and a moderate level of inhibitory potential was also observed. It is possible that these proteins, along with the associated RNA moieties, may not play a direct role in invasion. However, their interaction with parasite RNAs suggests that they may play some role(s) in ribosome assembly/function, immune modulation and cell-cell communication/signalling [52,53].

In summary, here we characterized two *P. falciparum* RAP domain proteins; PfRAP291 and PfRAP070 for parasite RNA binding and for their association with MSP-1 complex(s). The results show the binding of these proteins with *P. falciparum* RNAs in particular with 28s and 18s ribosomal RNAs associated with large and small ribosome subunits. We further show association of these proteins with some of the merozoite surface proteins as well, thus suggesting diverse role(s) of these proteins via mechanisms yet to be explored. Further knock-out or knock-down approaches will be required to know the exact role(s) of these proteins in *Plasmodium* biology of host parasite interaction.

## Declarations

### Ethical approval and consent to participate

All animal experiments conducted were approved by the Institutional Animals Ethics Committee of ICGEB (IAEC-ICGEB) under the approval no <u>ICGEB/AH/2015/01/MAL-74</u>. Liberian and Danish sera samples are collection of Dr Michael Theisen of Statens Serum Institut. Liberian samples used here were obtained in accordance with the Liberian Institute of Biomedical Research. Ethical approval for Danish blood donor samples was given by the Scientific Ethics Committee of Copenhagen and Frederiksberg, Denmark. Plasma samples from a total of 28 anonymous Danish blood, obtained for control purposes from Copenhagen University Hospital, were used. These individuals are resident of central Copenhagen and provided written consent to have a small portion of their blood stored, anonymously, and used for research purposes. Blood donors in Denmark were between the ages of 18 and 60. All data were analysed anonymously. Written informed consents were obtained from patients for all Indian sera samples (Ref. No. IESC/T-438/30.11.12). The use of sera samples in the study complied with the guidelines set by the Declaration of Helsinki.

### Consent for publication

Not applicable.

### Availability of data and materials

All data generated or analysed during this study are included in this published article (and its supplementary information files).

### Authors’ contributions

AA, AD, AP, IT, SP, SSS and NM performed research experiments; AA and RP did bioinformatics analysis; AA performed CLIP-seq experiments; AA, AD and ES performed inhibition assay; MT contributed tools; AA and PM designed the study, interpreted data and wrote manuscript; AM, DG and MT contributed to experiment design, data interpretation and manuscript preparation.

## Acknowledgements

AA acknowledges the financial support from University Grant Commission, Government of India in the form of UGC-JRF Fellowship. We thank Rotary blood bank, for providing human red blood cells. We also thank Prof. V. S. Chauhan (ICGEB, New Delhi) for providing MSP1-Fu clone. We acknowledge Bencos Research Solutions Pvt. Ltd., Thane, India, for their services in sequencing and CLIP-seq data analysis. We also thank Sheetal Kaul (Parasite Cell Biology Group, ICGEB, New Delhi) for her help in conducting CLIP-seq experiments and sequencing.

