## Supplementary Figures S1-S8 for "Characterization of *Plasmodium falciparum* RAP domain proteins-RAP291 and RAP070, and their association with ribosomal RNAs"

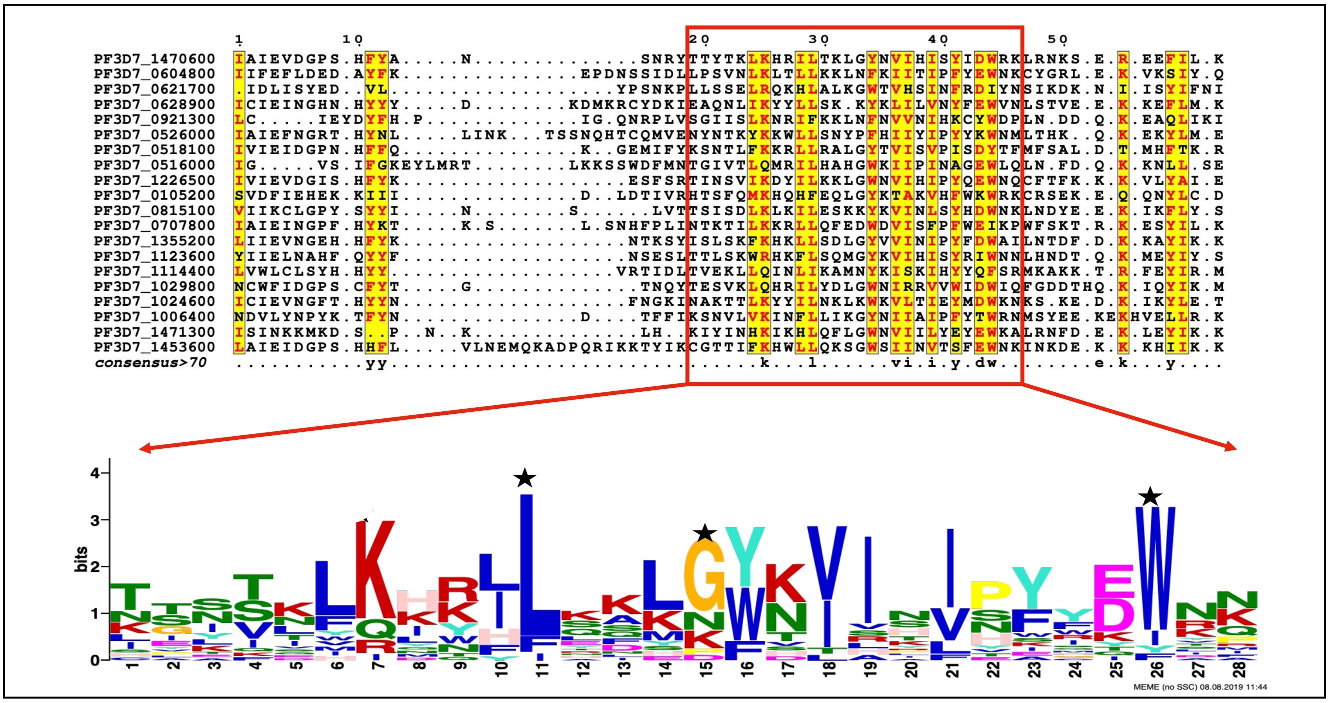


**Fig. S1-A.** Multiple Sequence Alignment of RAP domains of *Plasmodium falciparum* 3D7 RAP domain proteins. Motif was identified by MEME motif search tool. '*' star indicates the conserved residues predicted by Lee et al., 2006.

| Gene ID | start a.a. position | p value | sites |
|---|---|---|---|
| PF3D7_1470600 | 19 | 2.33E-22 | TTYTKLKHRILTKLGYNVIHISYIDWRK |
| PF3D7_1226500 | 19 | 2.17E-19 | TINSVIKDYILKKLGWNVIHIPYQEWNQ |
| PF3D7_1355200 | 19 | 1.43E-18 | ISLSKFKHKLLSDLGYVVINIPYFDWAI |
| PF3D7_0516000 | 25 | 1.61E-18 | TGIVTLQMRILHAHGWKIIPINAGEWLQ |
| PF3D7_1123600 | 19 | 2.95E-18 | TTLSKWRHKFLSQMGYKVIHISYRIWNN |
| PF3D7_0604800 | 23 | 6.45E-17 | LPSVNLKLTLLKKLNFKIITIPFYEWNK |
| PF3D7_1453600 | 33 | 1.23E-16 | CGTTIFKHWLLQKSGWSIINVTSFEWNK |
| PF3D7_1029800 | 19 | 2.88E-16 | TESVKLQHRILYDLGWNIRRVVWIDWIQ |
| PF3D7_1006400 | 19 | 1.63E-15 | KSNVLVKINFLLIKGYNIIAIPFYTWRN |
| PF3D7_0815100 | 19 | 1.63E-15 | TSISDLKLKILESKKYKVINLSYHDWNK |
| PF3D7_1024600 | 19 | 5.24E-15 | NAKTTLKYYILNKLKWKVLTIEYMDWKN |
| PF3D7_0518100 | 21 | 6.98E-15 | KSNTLFKKRLLRALGYTVISVPISDYTF |
| PF3D7_0105200 | 21 | 3.70E-14 | HTSFQMKHQHFEQLGYKTAKVHFWKWRK |
| PF3D7_0707800 | 24 | 5.30E-14 | NTKTILKKRLLQFEDWDVISFPFWEIKP |
| PF3D7_1471300 | 16 | 3.90E-13 | KIYINHKIKHLQFLGWNVIILYEYEWKA |
| PF3D7_0526000 | 30 | 4.61E-13 | NYNTKYKKWLLSNYPFHIIYIPYYKWNM |
| PF3D7_0921300 | 19 | 8.94E-13 | SGIISLKNRIFKKLNFNVVNIHKCYWDP |
| PF3D7_0621700 | 17 | 2.55E-12 | LLSSELRQKHLALKGWTVHSINFRDIYN |
| PF3D7_0628900 | 24 | 3.40E-11 | IEAQNLIKYYLLSKKYKLILVNYFEWVN |
| PF3D7_1114400 | 19 | 1.24E-10 | TVEKLLQINLIKAMNYKISKIHYYQFSR |

**FIG S1-B**. The Conserved Motif present in RAP domains of all *P. falciparum* RAP domain containing proteins with their p-values identified by MEME motif search tool(ver.5.0).

(i)

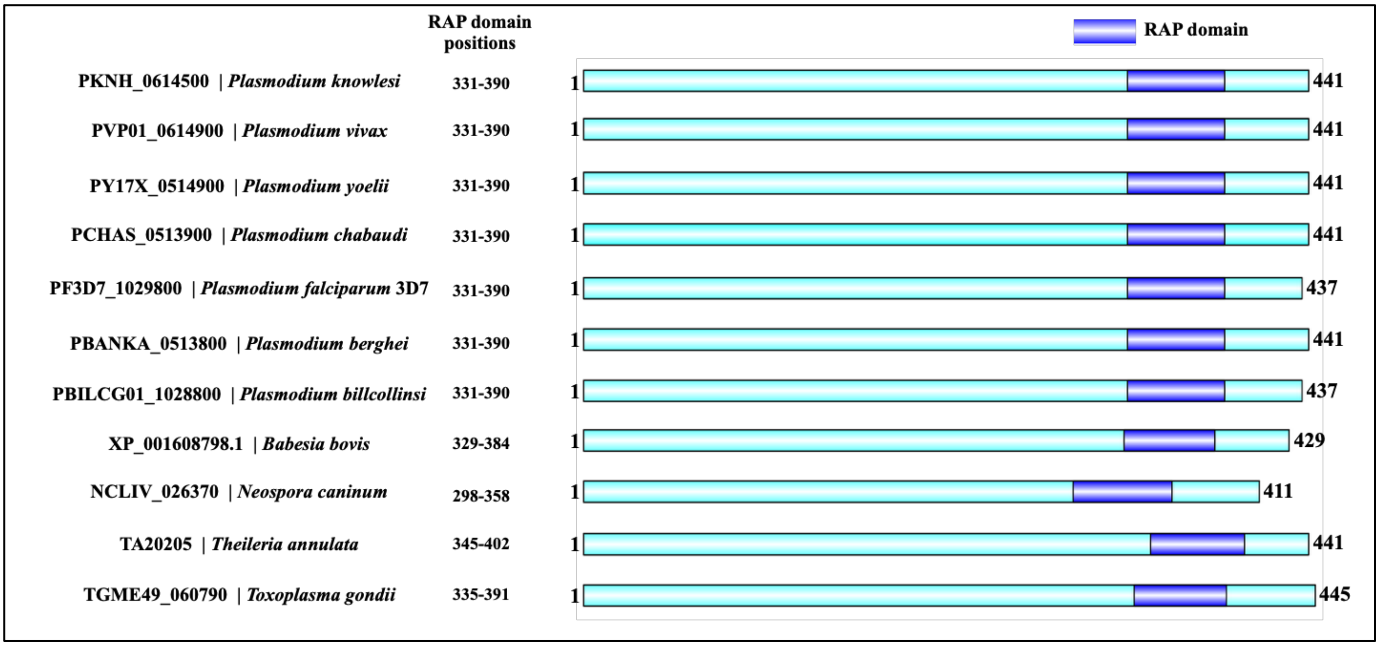


(ii)

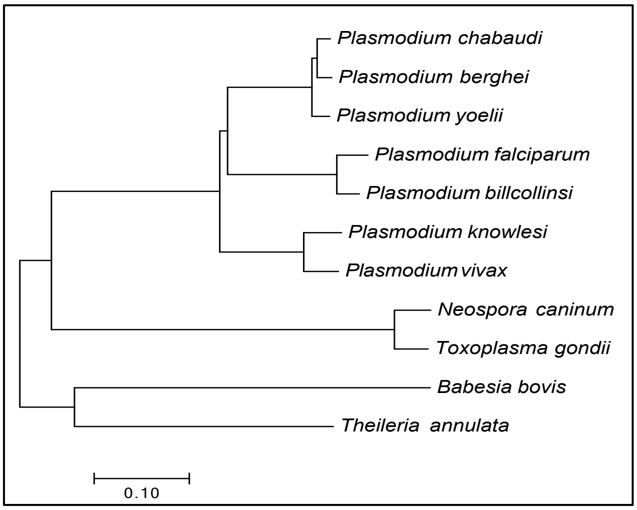


**FIG S1-C- (i)**. Schematics representation of the domain architecture of RAP291 (PF3D7_1029800) homologs across different apicomplexans. **(ii).** Phylogenetic analysis of RAP291 homologs from different apicomplexans.

(i)

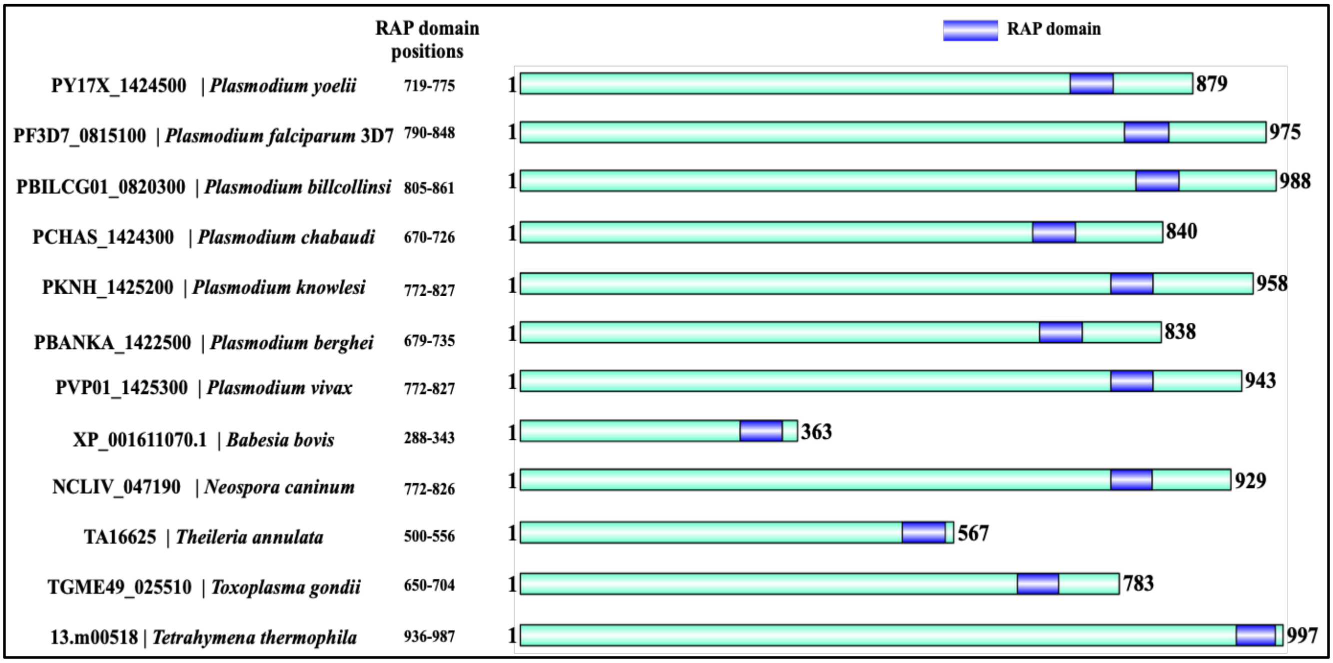


(ii)

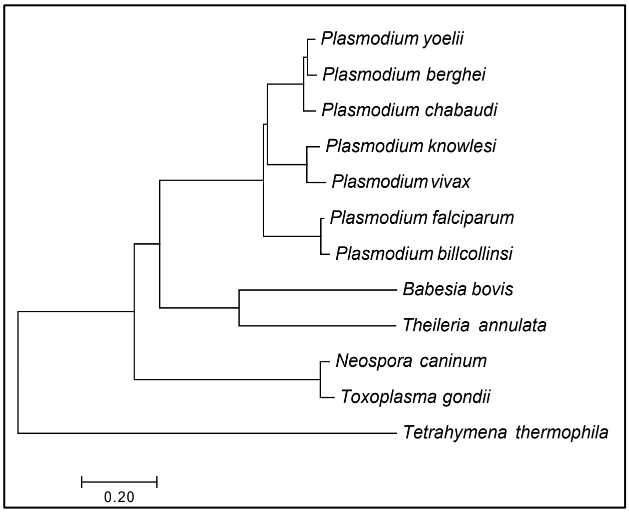


**FIG S1-D- (i)**. Schematics representation of the domain architecture of RAP070 (PF3D7_0815100) homologues across various apicomplexans and *T. thermophila*. **(ii)**. Phylogenetic analysis of RAP070 homologs from various apicomplexans and *T. thermophila*

**A**

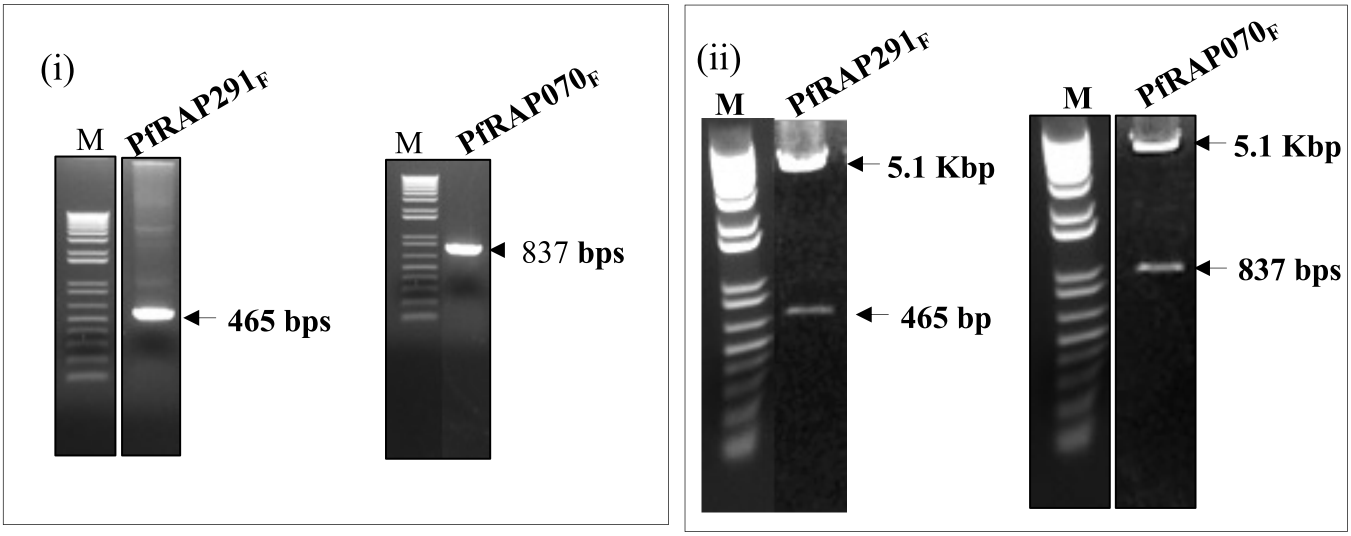


**FIG S2-A**. (**i**) PCR amplification of PfRAP291gene and PfRAP070 gene fragments. (**ii**) Restriction digestion analysis of the clones in pET28b expression vector.

**B**

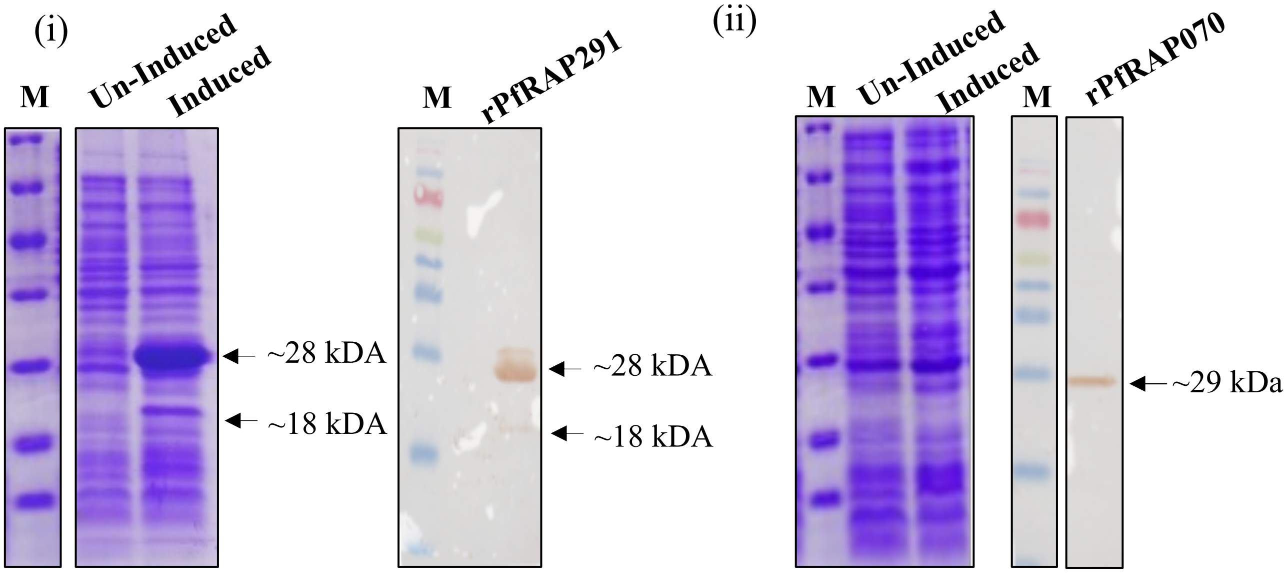


**FIG S2-B**. Coomassie and Western blot analysis to analyze the expression of rPfRAP291(**i**) and rPfRAP070(**ii**) in un-induced and induced *E. coli* cells.

(i)

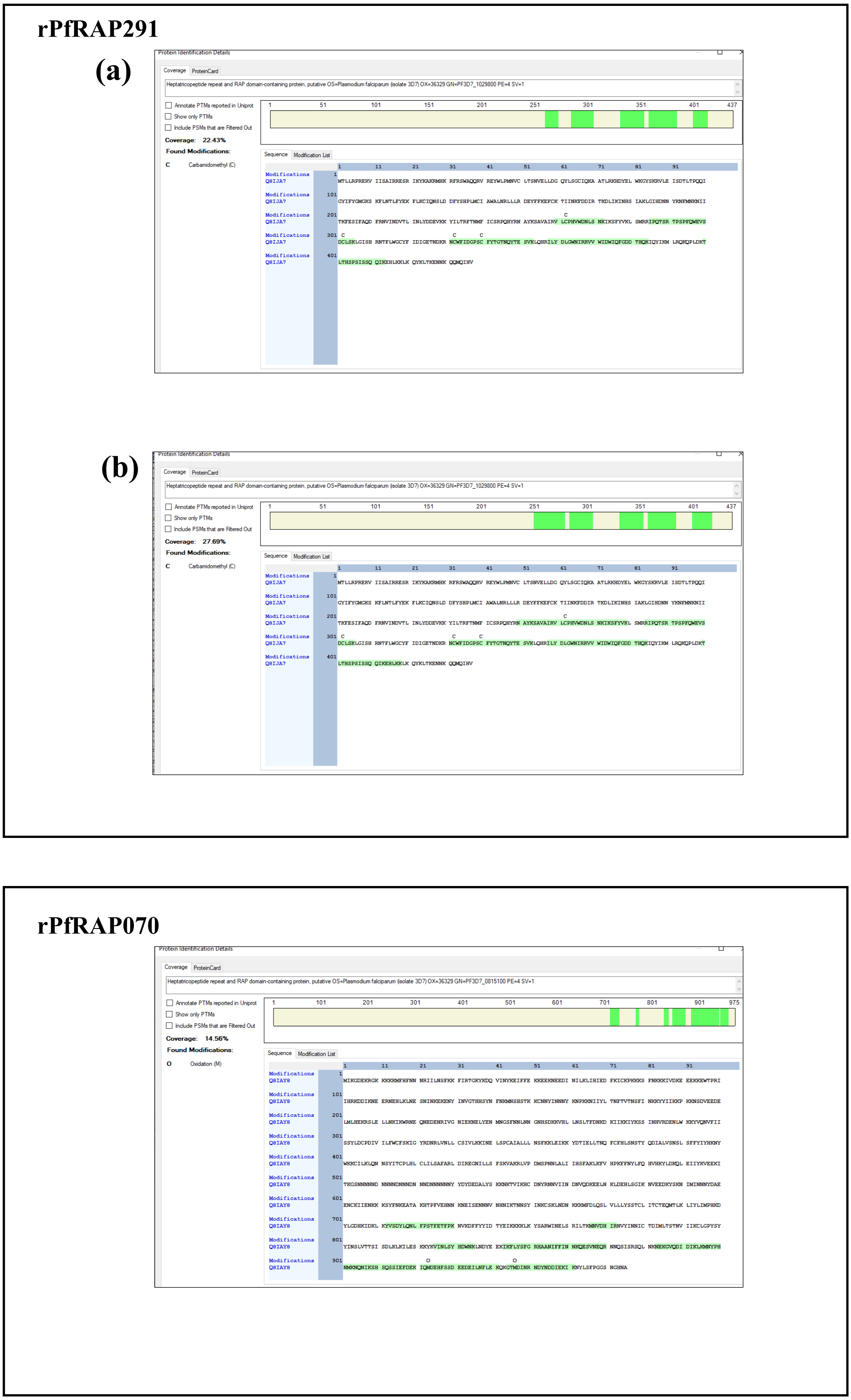


(ii)

**FIG S2-C**. Mass-spectrometry analysis data of the recombinant protein bands of rPfRAP291(**i-a**: for the higher molecular weight band, and **i-b**: for the lower molecular weight band) and rPfRAP070(**ii**) showing the coverage with the native proteins. The green highlighted regions indicate the peptides coverage area of the respective proteins, and the regions covered correctly indicate the recombinant proteins expression regions.

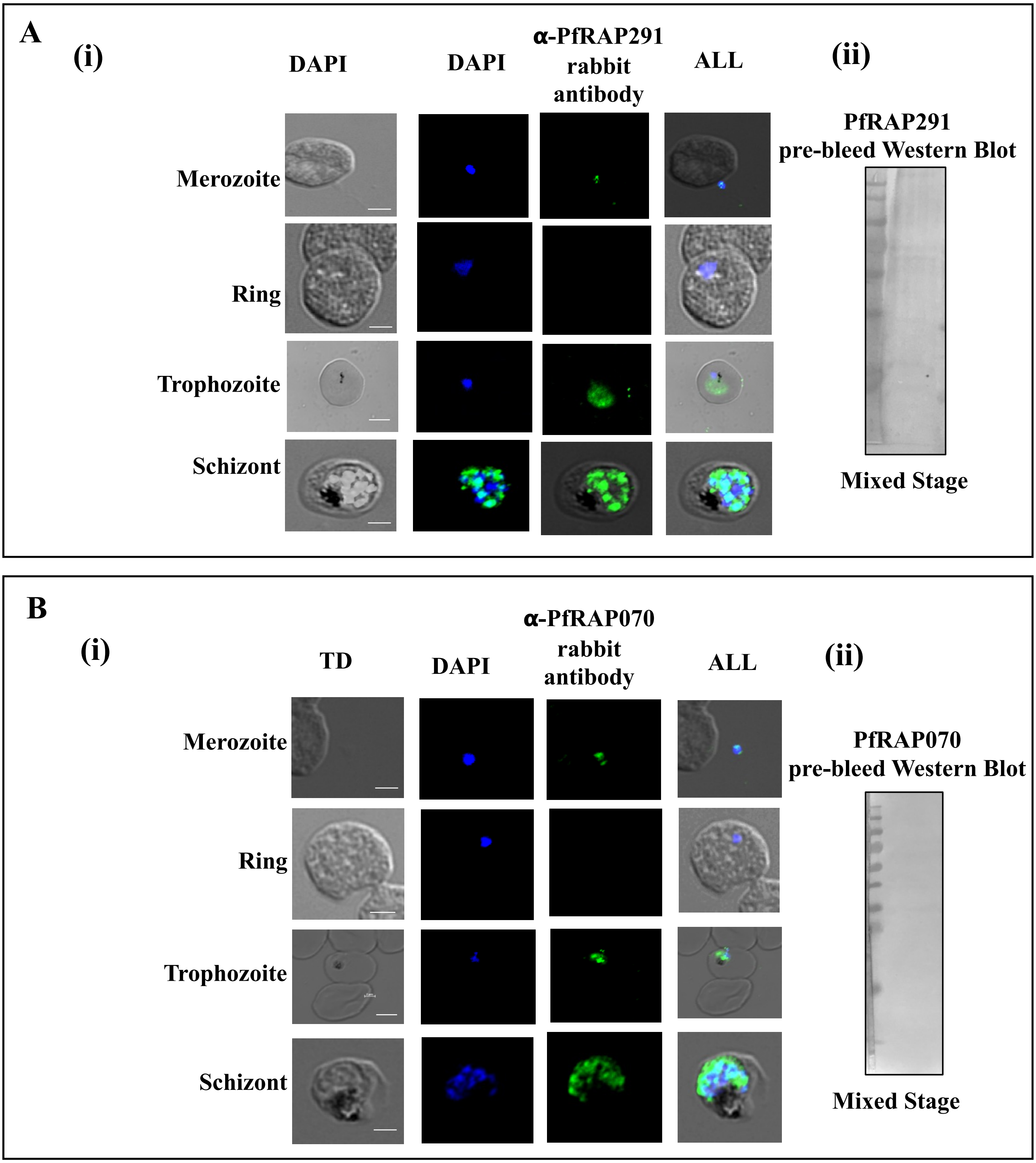


**FIG S3**. Localization of RAP291 and RAP070 in *P. falciparum* asexual blood stages. Confocal images showing immuno-staining in asexual blood stages of using **(A-i)** anti-PfRAP291 rabbit antibody and **(B-i)** anti-PfRAP070 rabbit antibody. A-ii and b-ii are Western blot analysis of mixed stage 3D7 parasite with rabbit pre-bleed of PfRAP291 and PfRAP070 respectively. Scale Bar represents 2 µm size.

(i) **PfRAP291**

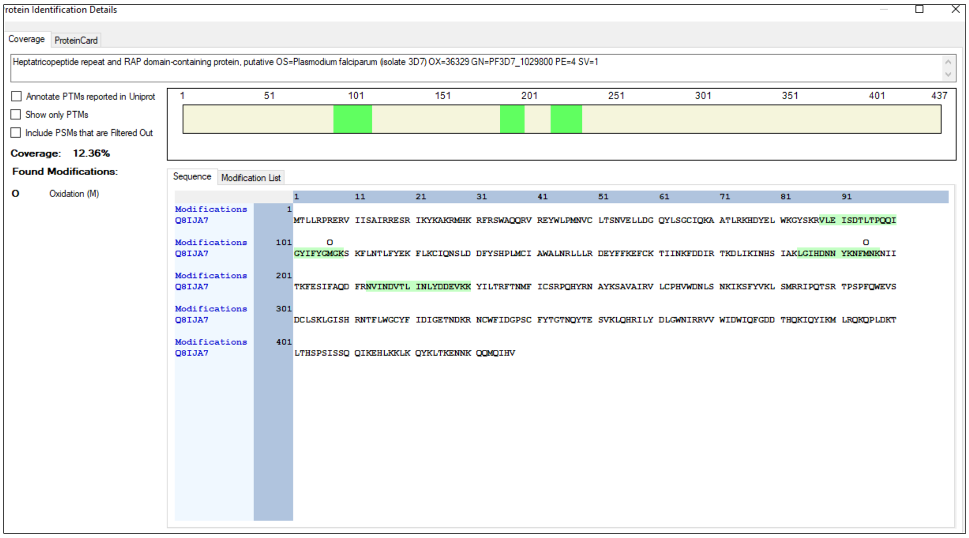


(ii) **PfRAP070**

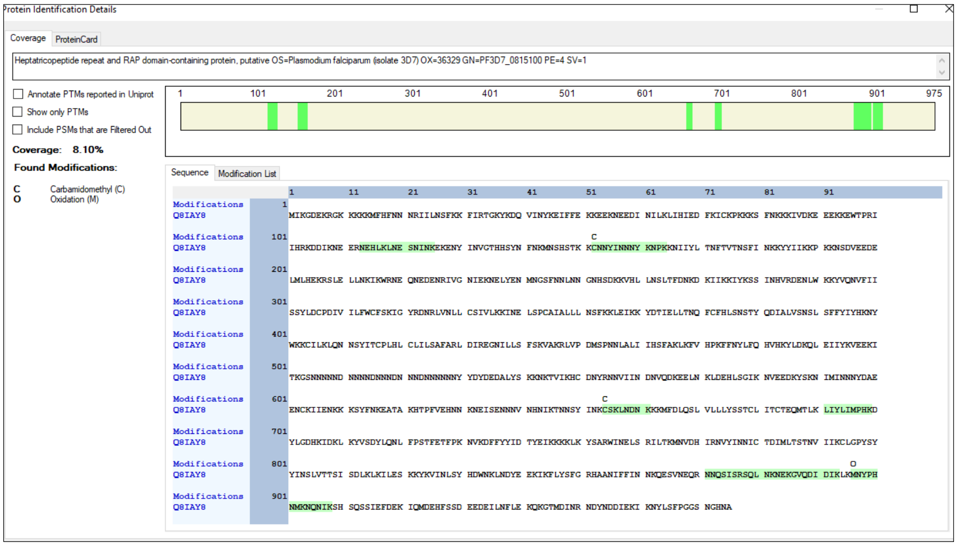


**FIG S3-C**. Mass-spectrometry analysis data of the native proteins bands in parasite lysate showing the coverage of the native RAP proteins (**i**-PfRAP291 and **ii**-PfRAP070) hence confirming the presence of the two RAP proteins at their respective band sizes.

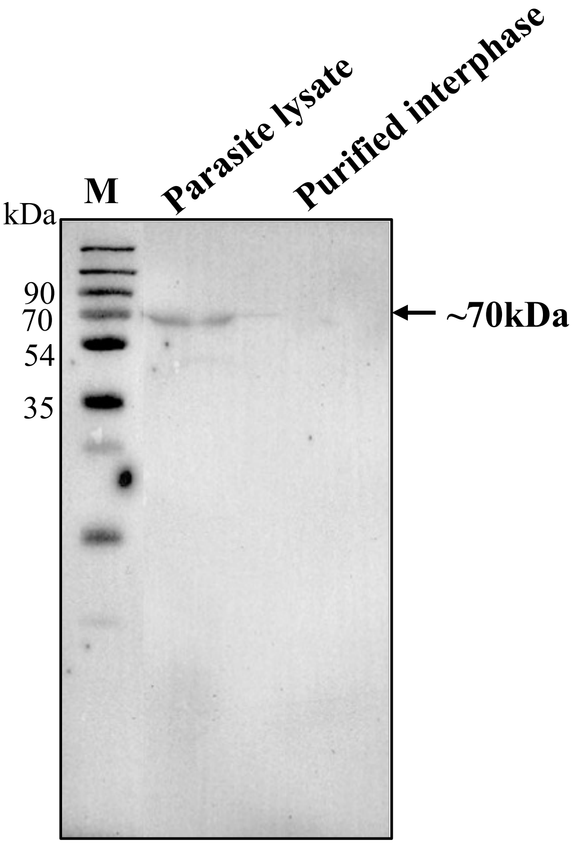


**FIG S4.** Western blot analysis showing the absence of ER resident protein, PfBiP, in purified interphase.

### Recombinant Proteins Used in Co-IP experiment

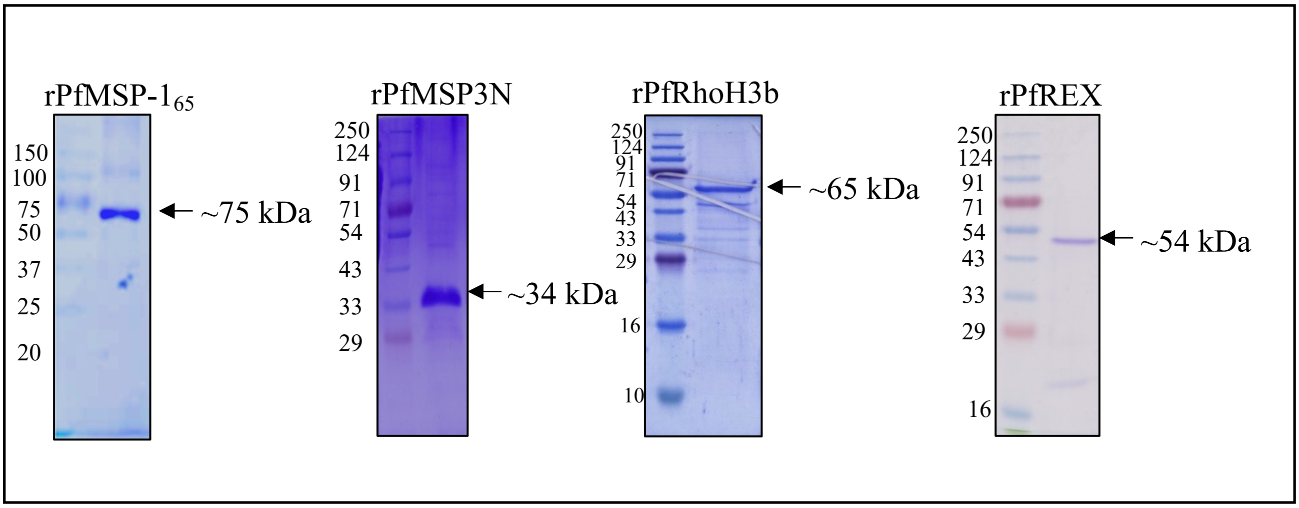


**FIG S5A.** Coomassie stained SDS PAGE to show the recombinant proteins used in Co-immunoprecipitation and Far-western analysis.

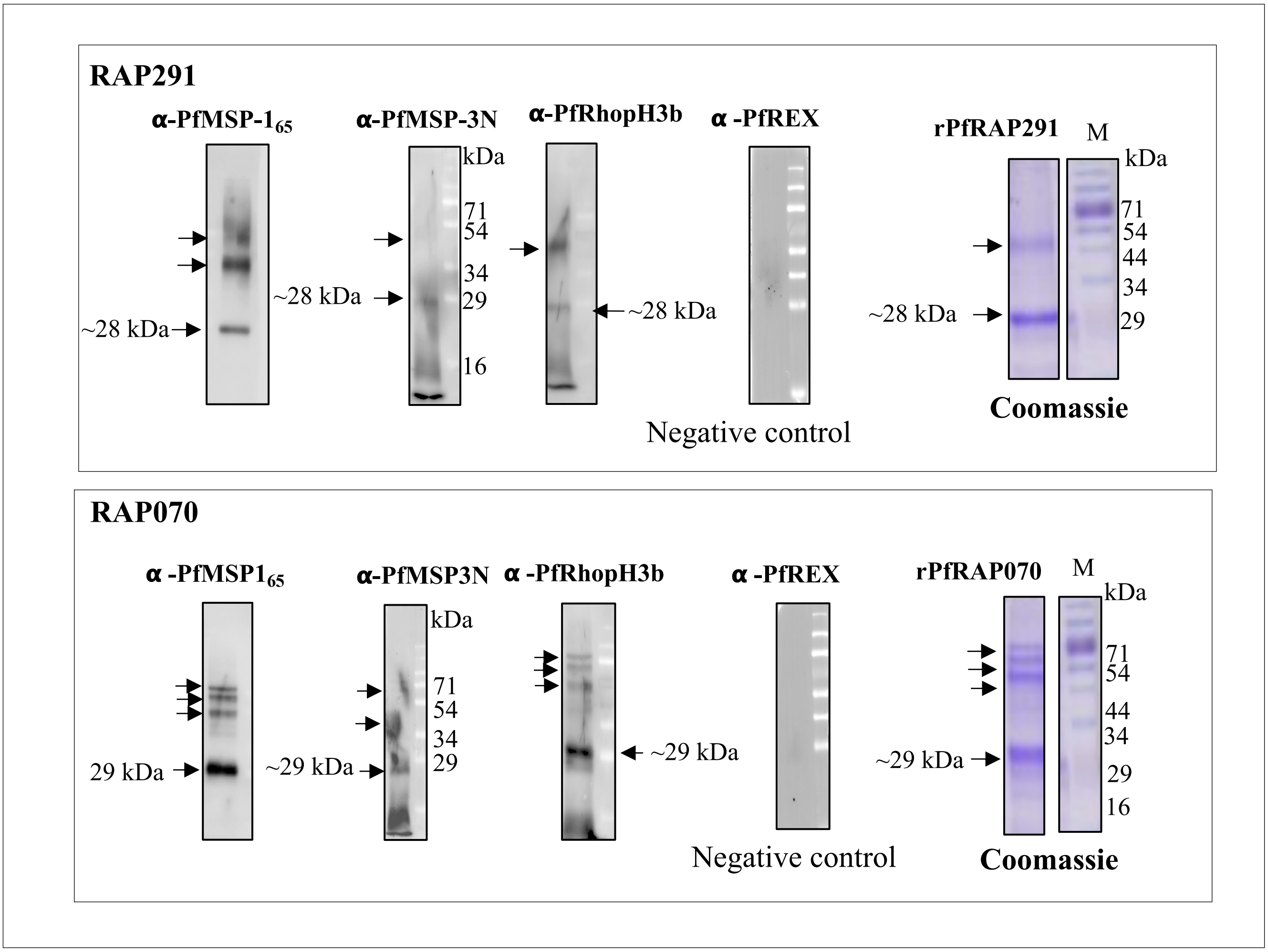


**FIG S5B.** Far-Western Analysis showing interactions of recombinant PfRAP291 and PfRAP070 (bait proteins) with rPfMSP-1$_{65}$, rPfMSP-3N, rPfRhopH3b all of these as prey proteins. PfREX protein did not show any interaction with either of the two the RAP domain proteins.

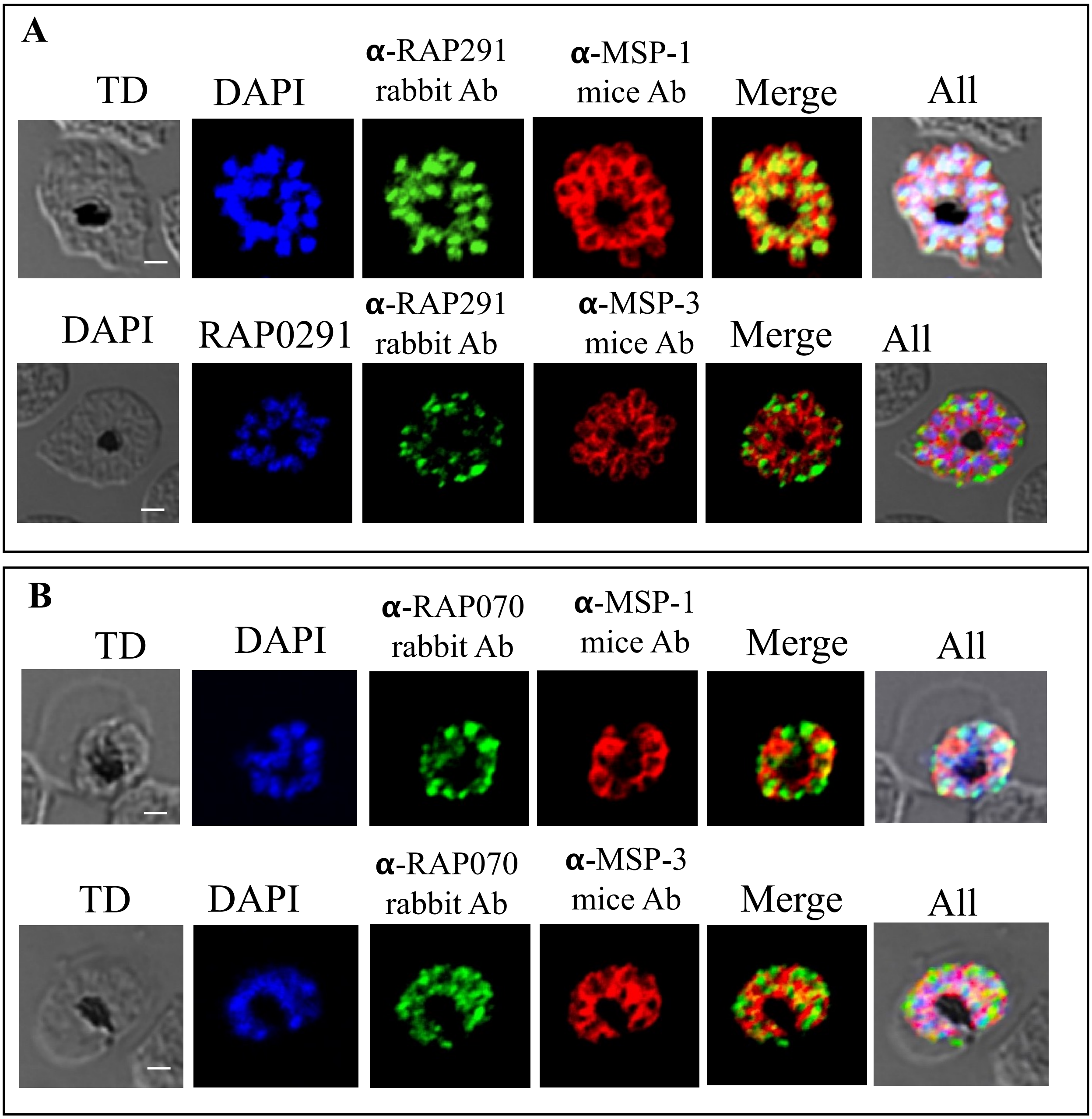


**FIG S6**. Co-localization of RAP proteins with MSP-1 and MSP-3 in *P. falciparum* schizont stage parasites. (**A)** Co-localization of PfRAP291 with MSP1 and MSP3. **B** Co-localization of PfRAP070 with MSP1 and MSP3. Scale bar represents 2 um size.

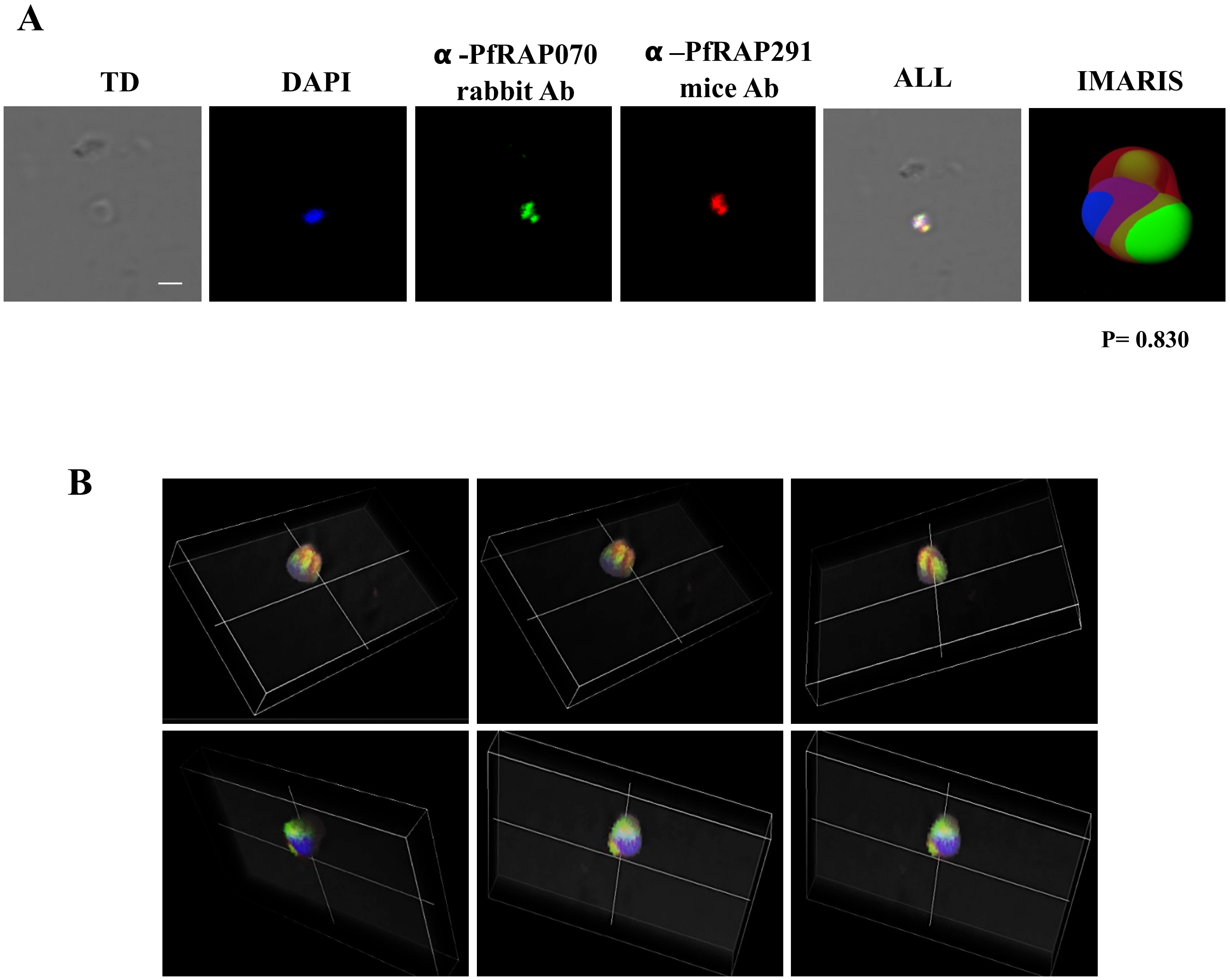


**FIG S7**. Co-localization Analysis of RAP291 with RAP070 proteins in *P. falciparum* merozoites. (**A**) Confocal images showing co-localization between rRAP070 vs rRAP291 at merozoite stage of the parasite using immunofluorescence assay. (P indicates Pearson's coefficient of co-localisation) (**B**) Movie screen shots generated to show 3 Dimensional overview of the co-localization. Scale bar represents 2 um size.

**RBC Binding Experiment**

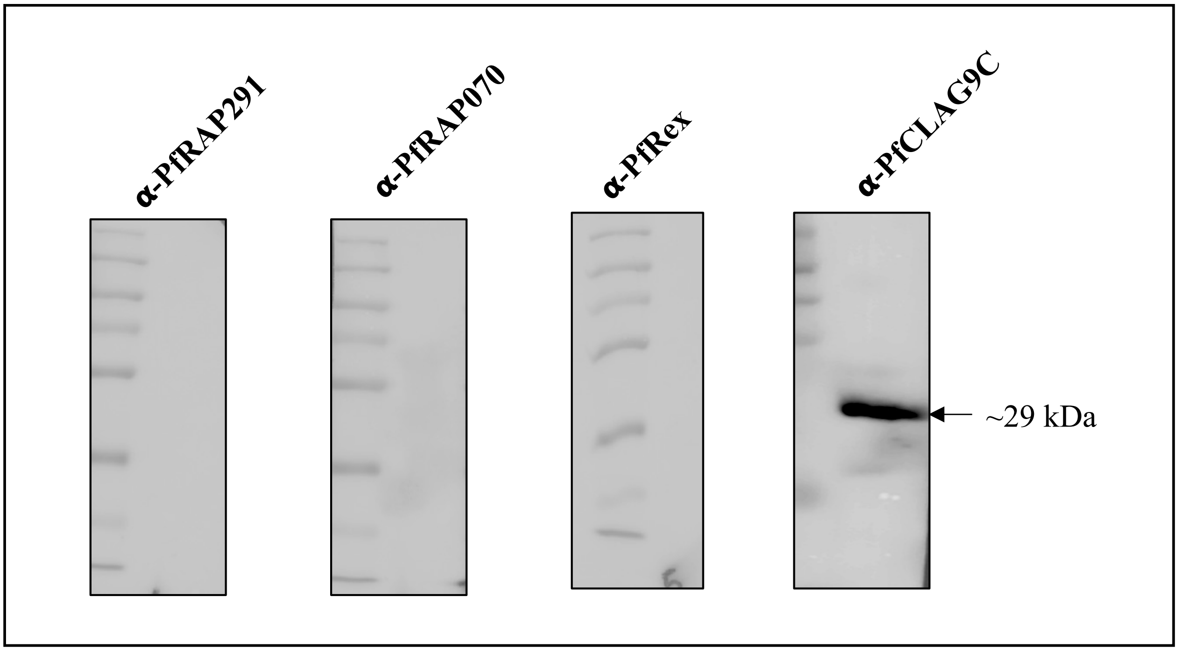


**FIG S8.** Recombinant PfRAP291 or PfRAP071 protein fragments do not bind human erythrocytes. rPfRAP291 and PfRAP070 proteins were incubated with uninfected human erythrocytes, and bound proteins were eluted from erythrocytes. Eluted proteins were analyzed on Western-blot using respective antibodies. PfREX was taken as a negative control and Clag9C was taken as a positive control.
